# Transcriptomic reprogramming screen identifies SRSF1 as rejuvenation factor

**DOI:** 10.1101/2023.11.13.566787

**Authors:** Alexandru M. Plesa, Sascha Jung, Helen H. Wang, Fawad Omar, Michael Shadpour, David Choy Buentello, Maria C. Perez-Matos, Naftali Horwitz, George Cai, Zhen-Kai Ngian, Carol V. de Magalhaes, Amy J. Wagers, William B. Mair, Antonio del Sol, George M. Church

**Affiliations:** Wyss Institute for Biologically Inspired Engineering at Harvard University, Boston, MA, USA; Department of Genetics, Harvard Medical School, Boston, MA, USA; Computational Biology Group, CIC bioGUNE-BRTA (Basque Research and Technology Alliance), Bizkaia Technology Park, Derio, Spain; Department of Biological Engineering, MIT, Cambridge, MA, USA; Paul F. Glenn Center for the Biology of Aging, Harvard Medical School, Boston, MA, USA; Department of Molecular Metabolism, Harvard T.H. Chan School of Public Health, Harvard University, Boston, MA, USA; Department of Stem Cell and Regenerative Biology, Harvard University, Cambridge, MA, 02138 USA; Computational Biology Group, Luxembourg Centre for Systems Biomedicine (LCSB), University of Luxembourg, L-4362 Esch-sur-Alzette, Luxembourg; Ikerbasque, Basque Foundation for Science, Bilbao, Bizkaia, 48012, Spain

## Abstract

Aging is a complex process that manifests through the time-dependent functional decline of a biological system. Age-related changes in epigenetic and transcriptomic profiles have been successfully used to measure the aging process^1,2^. Moreover, modulating gene regulatory networks through interventions such as the induction of the Yamanaka factors has been shown to reverse aging signatures and improve cell function^3,4^. However, this intervention has safety and efficacy limitations for *in vivo* rejuvenation^5,6^, underscoring the need for identifying novel age reversal factors. Here, we discovered SRSF1 as a new rejuvenation factor that can improve cellular function *in vitro* and *in vivo*. Using a cDNA overexpression screen with a transcriptomic readout we identified that SRSF1 induction reprograms the cell transcriptome towards a younger state. Furthermore, we observed beneficial changes in senescence, proteasome function, collagen production, and ROS stress upon SRSF1 overexpression. Lastly, we showed that SRSF1 can improve wound healing *in vitro* and *in vivo* and is linked to organismal longevity. Our study provides a proof of concept for using transcriptomic reprogramming screens in the discovery of age reversal interventions and identifies SRSF1 as a promising target for cellular rejuvenation.

## Introduction

Aging is a highly complex process modulated by evolutionarily conserved genetic pathways. Across generations, aging is reversed through a replay of the developmental program, and it has been proposed that the right signals can induce a similar age reversal in somatic cells of adult organisms^7,8^. In agreement with this hypothesis, it was shown that the induction of the four Yamanaka factors (OCT4, SOX2, KLF4, and MYC, or OSKM) not only reprograms aged somatic cells to an induced pluripotent stem cell (iPSC) state^9^, but also reverses several features of aging such as cellular senescence, telomere size, oxidative stress, mitochondrial metabolism, and transcriptomic and epigenetic profiles^4,10^. Moreover, several recent studies have reported rejuvenating effects by partially recapitulating the reprogramming process *in vivo*^11–15^.

Nonetheless, OSKM induction poses significant neoplastic risk due to loss of cell identity, and it has varying degrees of efficiency across different tissues^3,5,6,16,17^. However, it is possible to achieve age reversal without dedifferentiation^7,18^, and a recent study showed that there are several sets of factors that can perform cellular rejuvenation with varying degrees of cell identity loss^19^. Therefore, a promising future direction for the aging field is the search for novel, safe, and effective factors that can reverse the age of human tissues with high specificity and without losing cell identity.

An efficient approach for uncovering such targets is to focus on rejuvenating the transcriptome, which is a key determinant of cell identity and function. With age, there are predictable changes in gene expression^20,21^, which serve as the basis for aging assays at the epigenetic and transcriptomic levels^1,2^. Moreover, there is a growing body of evidence that supports a systems-level view of cell engineering, whereby exogenous signals in the form of genetic perturbations can induce desired cell states^19,22,23^. Thus, the transcriptome is an important and accessible layer of gene regulation that can be used to read and write cell states, including age related changes^24,25^.

In this study, we sought to discover novel age reversal factors by using a cDNA overexpression screen to identify genes that can reprogram the transcriptome of old cells to a younger state. We first developed a transcriptomic aging assay that can accurately measure the age of human fibroblasts and respond to known biological interventions. Then, we screened candidate genes for their ability to restore a young transcriptome in old cells. We uncovered that these cell state conversions were influenced by the identity, dosage, and timing of our perturbation, and also by the donor-specific aging phenotypes of our fibroblasts. Our screen discovered SRSF1 as a novel age reversal factor, whose overexpression reprogrammed the cellular transcriptome to a more youthful state. The rejuvenation effect was driven by gene expression changes in the histone methylation and translation modules. Finally, we showed that SRSF1 induction rejuvenated several features of cellular aging and reversed the wound healing dysfunction of aged mice.

### Measuring aging using gene expression

To develop a transcriptomic aging assay, we trained a machine learning predictor using RNA-sequencing (RNA-Seq) data from primary normal human dermal fibroblasts (NHDFs) aged 1-96 years^2^. Our approach focused on functional interpretability by sub-setting the transcriptome into different cellular processes and searching for those most predictive of chronological age (Fig. 1a). This approach stably selected eight processes (WNT signaling, Actin polymerization, Junction organization, Histone methylation, ER quality control, Translation initiation, Sodium transport, and Lipid transport) with varying contributions to the model (Fig. 1b, Extended Data Fig. 1a, b). We used a linear combination of the normalized expression values of a subset of genes (146) involved in these eight processes to generate an accurate transcriptomic aging assay for human fibroblasts (Extended Data Fig. 1c, Extended Data Table 1). When testing our predictor on 12 different fibroblast datasets (Fig. 1c), we observed a comparable performance to that of DNA methylation (DNAm) predictors^26^ (Pearson’s r=0.93, p-value < 2.2e-16; MAE=6.66). Moreover, when studying the gene expression changes of the eight cellular processes across aging, we identified three distinct trajectories (Extended Data Fig. 1d) with histone methylation as the earliest process to decline with age, which aligns with recent data suggesting epigenetic changes play a causal role in the aging process^27^.

**Figure 1.**
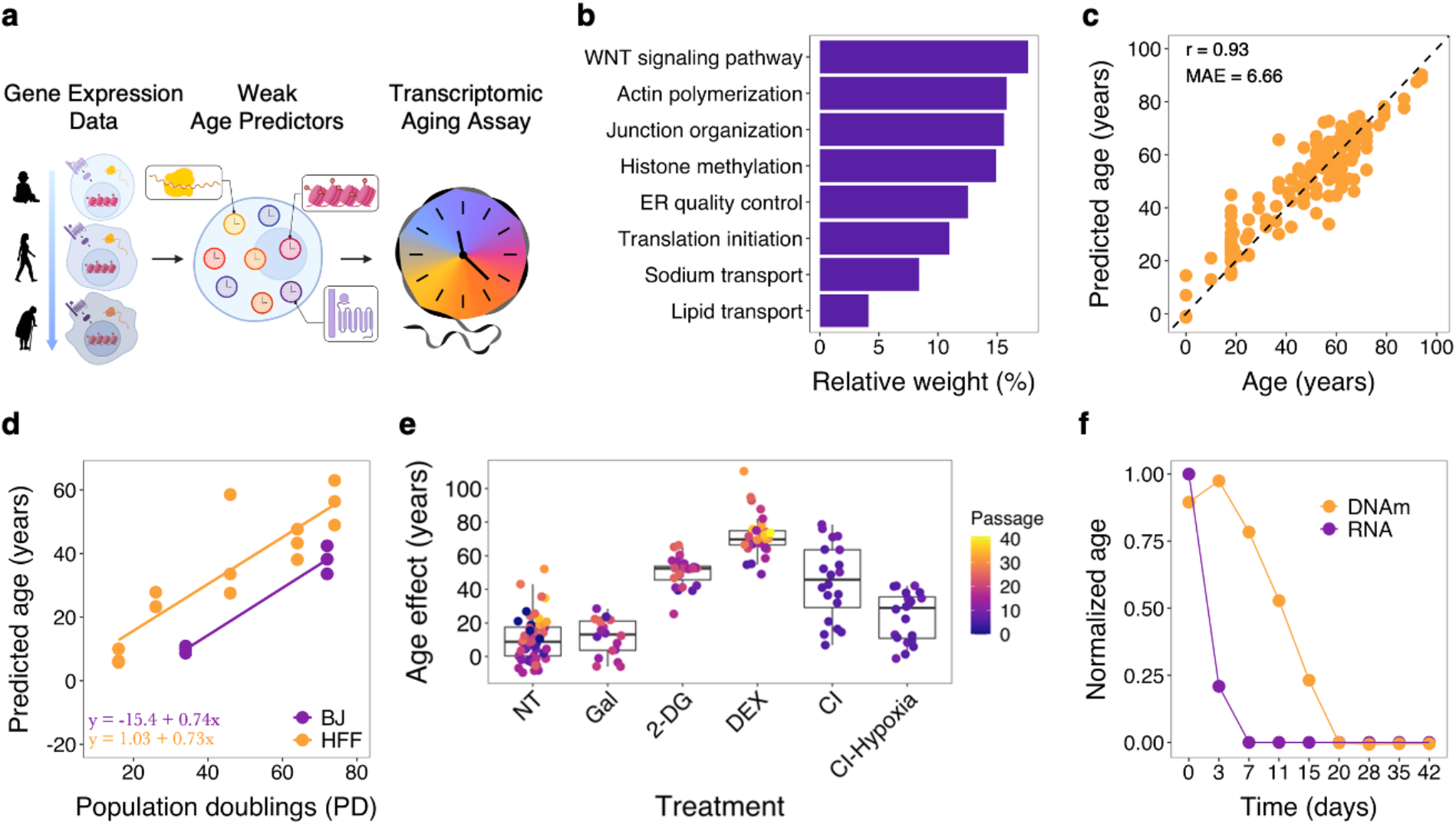
Gene expression is an accurate and robust aging biomarker a, Schematic for developing the transcriptomic aging assay. Gene expression data from human fibroblasts of different ages was used to train process-specific weak age predictors, which were combined in a sparse linear model to select the most predictive combination thereof. The expression levels of 146 genes involved in the most predictive processes were the inputs for the transcriptomic aging assay. **b**, Relative weights of the eight processes part of the transcriptomic assay. **c,** Predictive accuracy of our transcriptomic assay applied to 150 fibroblast samples from 12 different public datasets. **d,** Aging rates of cultured NHDFs lines as determined by our transcriptomic predictor. BJ, HFF = foreskin fibroblast lines. **e,** Response of transcriptomic assay to different *in vitro* stressors. NT = non-treated, Gal = galactose, 2-DG = 2-deoxy-D-glucose; DEX = dexamethasone; CI = contact inhibition; hypoxia = 3% O_2_. **f,** Predicted age (normalized to day 0) of fibroblast reprogrammed with OSKM using our transcriptomic assay and a DNAm clock^26^.

Next, we tested our assay’s responsiveness to known age-modulating interventions, to confirm that we can use gene expression as a proxy measurement of cell function. It was previously shown, using the DNAm clock^28^, that NHDFs age upon *in vitro* culturing at a rate of 1 year/0.7 population doublings (PD), and our transcriptomic assay confirmed this observation in two different skin fibroblast lines (Fig. 1d). Moreover, we showed that lung fibroblasts have different aging rates (Extended Data Fig. 1e) and the transcriptomic model predicted pluripotent stem cells to be significantly younger than fibroblasts (Extended Data Fig. 1f). We also confirmed an age increase in fibroblasts subjected to senescence-inducing FOXM1 knock-down (Extended Data Fig. 2a), treated with cellular stressors^29^ (Fig. 1e), exposed to UV-damage (Extended Data Fig. 2b), collected from progeroid donors (Extended Data Fig. 2c), and immunostimulated with polyinosinic -polycytidylic acid (poly(I:C)) (Extended Data Fig. 2d). Importantly, our transcriptomic assay also captured the age reversal effects of cellular reprogramming (Fig. 1f) and Rapamycin (Rapa) treatment (Extended Data Fig. 2e-f), confirming our ability to detect both pro- and anti- aging gene expression changes, which we failed to observe using another published RNA aging assay (Extended Data Fig. 2 g-k)^30^. Interestingly, in comparison to a commonly used epigenetic clock, our transcriptomic assay showed a faster response to both Rapa and OSKM (Fig. 1f, Extended Data Fig. 2f), reflecting the more dynamic nature of gene expression compared to DNAm.

### Screening for age reversal genes

To identify novel age reversal factors, we conducted a transcriptomic reprogramming screen^25^. First, we generated a list of candidate age-modulating genes using a network scoring approach^31^ (Extended Data Fig. 3a). We chose the top 89 genes (Extended Data Table 2, Extended Data Fig. 3b, c), along with previously reported positive (pro-aging: KAT7^32^, CXCR2^33^; anti-aging: ATG5^34^, HES1^35^, OSKM^4^) and negative (blue fluorescent protein (BFP)) controls, to screen for their ability to reprogram the transcriptome of old NHDFs to a younger state. Next, we generated stable overexpression lines in aged fibroblasts from three different aged donors (M55, M65, M79), which we then used to induce the candidate genes for three days and measure age-related changes through RNA-Seq and flow cytometry-based assays (Fig. 2a). Our data showed a wide range of gene induction levels, ranging from no overexpression to a 4000-fold increase (Fig. 2b). However, the overexpression levels were consistent between the three NHDF lines, indicating that induction differences were gene specific. We observed that fold change levels were inversely correlated with the endogenous expression level for each gene (Extended Data Fig. 3d), confirming our previous findings of a transcriptional output limit^31^.

**Figure 2.**
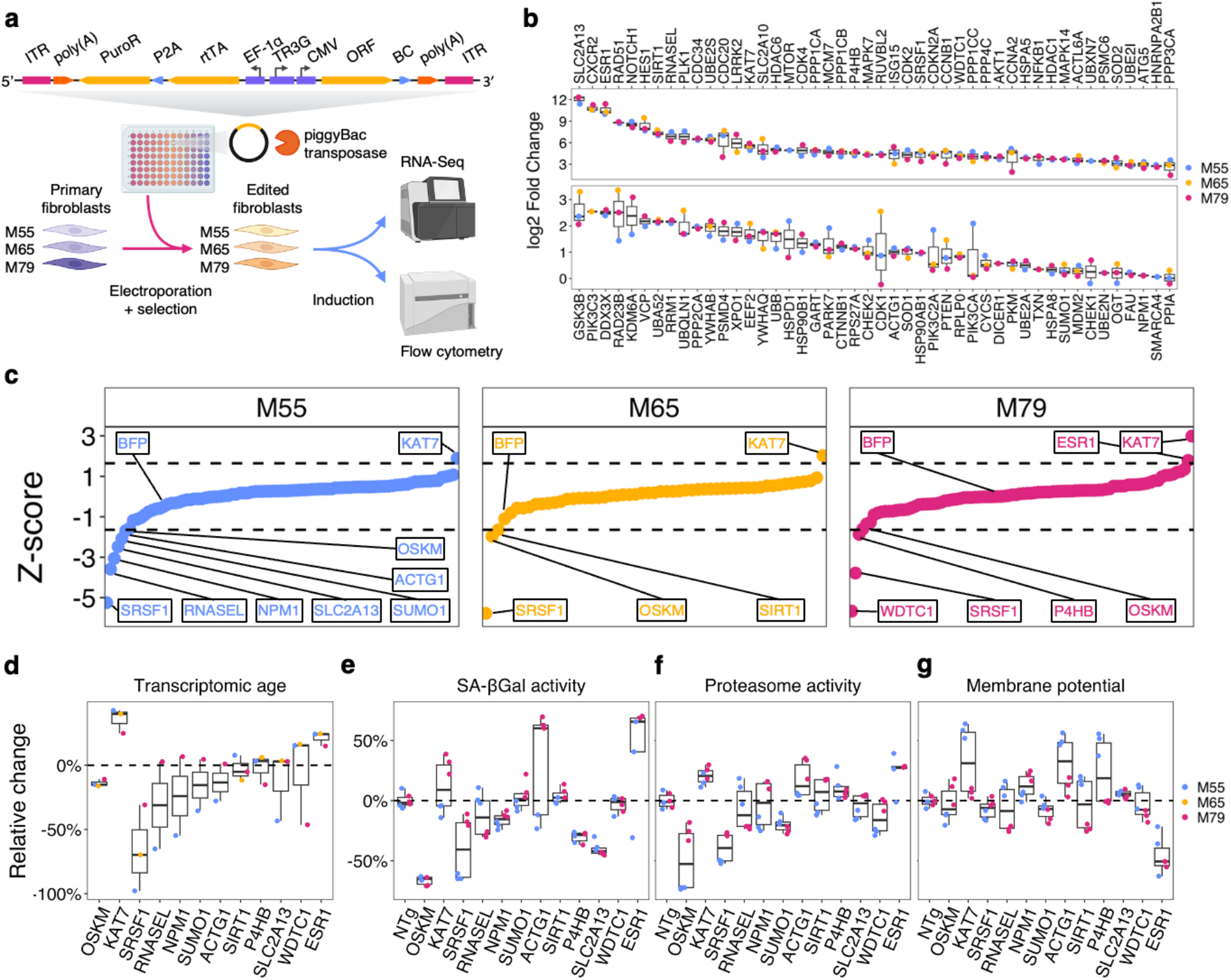
cDNA overexpression screen identifies age-modulating genes a, Workflow of cDNA overexpression screen for rejuvenating interventions. We electroporated 95 different plasmids containing our gene of interest in an overexpression cassette along with a piggyBac transposase expressing plasmid into three different NHDF lines. The edited fibroblasts were induced using Dox (1 ug/mL) for three days and assayed for age-related changes using RNA-Seq and flow cytometry assays. **b,** Heatmap of gene induction for all our single-gene overexpression lines in the three different NHDF backgrounds (n=2/line). There is a consensus across the log2 FoldChange levels across the three different lines, and we observed a range of induction values across all tested genes. **c,** Z-score for the age effect measured by our transcriptomic assay in the three NHDF lines (n=2). Genes with a significant effect on the transcriptomic age (|z-score| > 1.65 for 0.05 significance level) and BFP are labeled. **d,** Relative change in the predicted age of the cell lines upon gene induction for all our hits (|z-score| > 1.65). **e-g,** Staining data for representative gene hits and controls across the three assayed cellular processes in the M55 and M79 lines (n=3). Data is normalized with respect to the gene-specific No Doxycycline control and the line specific gene control.

Then, we analyzed the effect of each genetic perturbation on the aging phenotype of the three NHDF lines using our transcriptomic assay. Our predictor correctly estimated the ages of the WT NHDF lines (Extended Data Table 3) and was able to detect age-modulating effects for some of our controls (Fig. 2c). As expected, OSKM had a strong rejuvenating effect across all three lines, while KAT7 had the highest pro-aging effect. Similar to our BFP control, ATG5, HES1, and CXCR2 did not show any significant effects on our transcriptomic assay, despite achieving robust overexpression levels. This could indicate that, for some genes, three days of induction is not sufficient to observe age-related effects at the gene expression level using our assay. Nevertheless, we discovered nine genes (SRSF1, RNASEL, NPM1, SUMO1, ACTG1, SIRT1, P4HB, SLC2A13 and WDTC1) that showed a significant age reversal in at least one NHDF line, and one gene (ESR1) that robustly increased the transcriptomic age across all three lines (Fig. 2d).

SRSF1 is a splicing factor whose expression decreases in old age^36^ and is associated with parental longevity in humans^37^. RNASEL is an interferon-activated endoribonuclease whose levels in human serum are inversely correlated with metabolic syndrome and age^38^. NPM1 is involved in ribosomal biogenesis and was recently identified as a regulator of HSC aging^39^. SUMO1 (ubiquitin-like protein involved in protein stability), ACTG1 (structural protein part of the cellular cytoskeleton), P4HB (abundant multifunctional enzyme), and WDTC1 (protein of unknown function) currently have no reported link to aging phenotypes. SIRT1 is a histone deacetylase whose role in aging has been thoroughly studied^40^. SLC2A13 is a proton myo-inositol cotransporter that decrease in expression during aging in the human dorsolateral prefrontal cortex^41^ and was recently identified as a risk gene for Parkinson’s disease^42^. Lastly, ESR1 is a nuclear receptor for the estrogen hormone with connections to Alzheimer’s^43^.

For orthogonal readouts of cell function, we also performed three staining assays that report on important cellular features: senescence associated β-galactosidase activity (SA-βGal activity), proteasomal activity, and mitochondrial membrane potential in two NHDF lines (M55 and M79) (Extended Data Fig. 4). Our data showed that OSKM, SRSF1, NPM1, SLC2A13, and P4HB reduced the senescence phenotype in both donor lines, while KAT7 and ESR1 increased it (Fig. 2e). Surprisingly, the proteasome assay showed decreased activity in the OSKM, SRSF1, and SUMO1 overexpression lines but increased activity in the KAT7, ACTG1, P4HB, and ESR1 fibroblast lines (Fig. 2f). These effects on the 20S proteasome activity were unexpected, but they were consistent between the pro-aging (KAT7 and ESR1) and anti-aging (OSKM and SRSF1) interventions, suggesting decreased necessity of proteasomal activity and potential increased proteostasis^44^. In the mitochondrial assay, KAT7, NPM1, ACTG1, and SLC2A13 increased the membrane potential, while ESR1 drastically decreased it. Interestingly, SIRT1 had significant diverging effects, increasing the mitochondrial potential of M55 and decreasing it in M79. This, along with our prior observations of large variability in mitochondrial membrane potential across different donors, suggests the assay is susceptible to biological noise. Finally, the observed differential responses between the NHDF lines in both our transcriptomic and flow cytometry assays (Fig. 2d-g, Extended Data Fig. 4) indicate that donor-specific differences in the initial transcriptomic state might influence the outcome of our perturbations.

### Reprogramming the transcriptome of aged fibroblasts

To validate our age reversal screen hits, we generated new overexpression lines of SRSF1, SIRT1, WDTC1, RNASEL, and SLC2A13 in six aged NHDFs (M55, M65, M67, M68, M69, M79). Then, we induced the transgenes and applied our transcriptomic aging assay. Notably, we confirmed the observed age reversal effect of SRSF1 in all six lines, with an average rejuvenation of ∼26 years (Fig. 3a). SIRT1 and RNASEL also had rejuvenating effects of varying degree in four out of six lines (Fig. 3a), while WDTC1 and SLC2A13 no longer showed significant beneficial effects.

**Figure 3.**
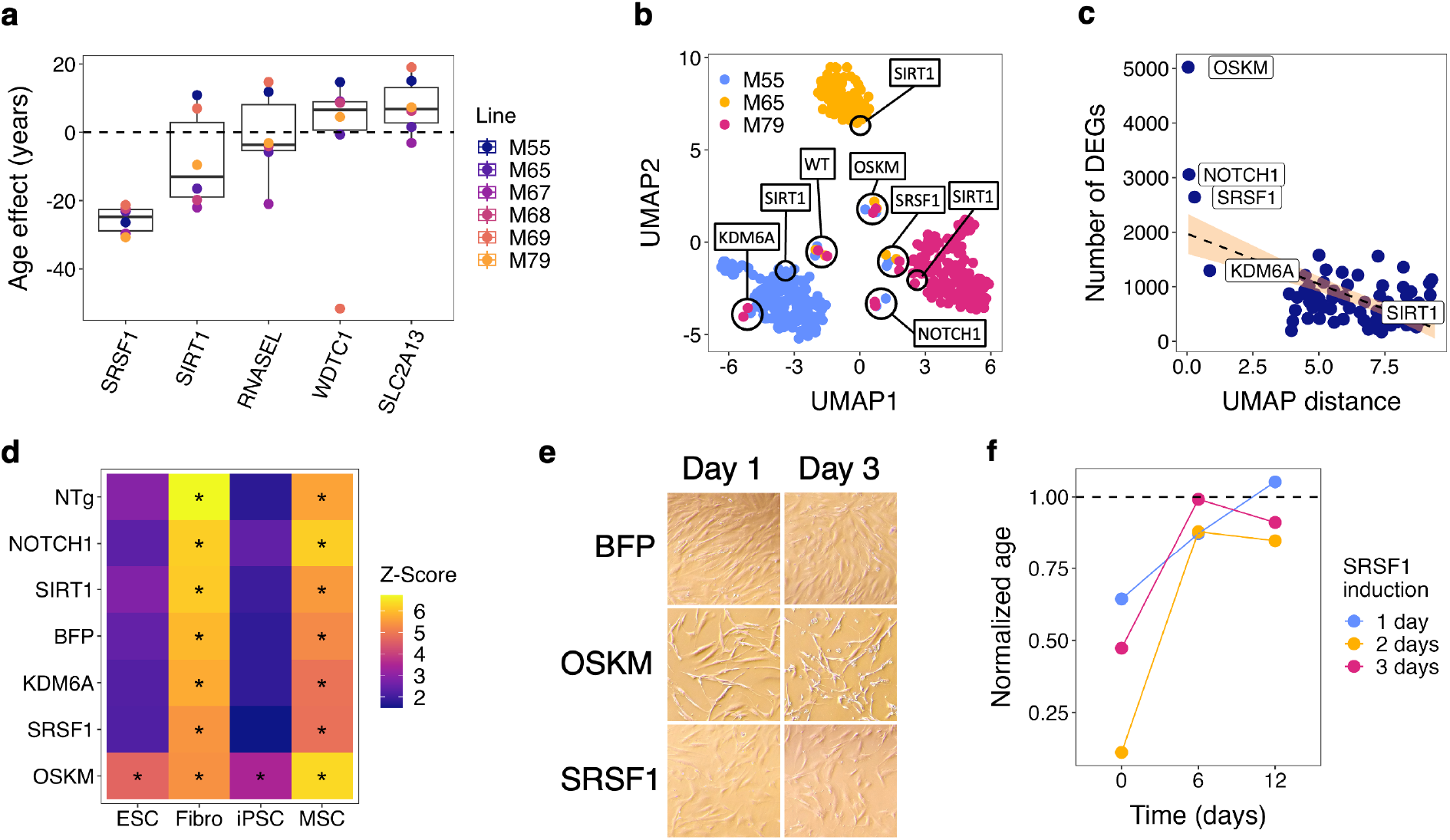
Cell state and perturbation influence transcriptomic reprogramming outcomes a, Age effect (predicted age – age) of hit overexpression in validation experiments using six NHDF lines (n=2/line). We observed robust age reduction in the SRSF1 samples, and a beneficial effect of SIRT1 on the transcriptome in four out of six lines. The other three genes had weak or cell-line specific effects. **b,** Total transcriptome UMAP of overexpression lines from age reversal screen. There is a strong clustering based on donor cell lines with only a few perturbations overcoming line-specific differences. **c,** Relationship between transcriptomic variability of interventions across cell lines and their induced transcriptomic changes, measured as number of DEGs. We observed a strong inverse correlation (Pearson’s r=-0.486; p-value=1.39e-06) between the mean UMAP distance of biological replicates of the same gene and the associated number of DEGs, with OSKM, NOTCH1, SRSF1, and KDM6A having the largest effects. **d,** Enrichment analysis for gained identities with respect to a fibroblast transcriptome. Data is averaged across the three NHDF lines (M55, M65, M79) and shows that only OSKM-expressing cells significantly acquire ESC and iPSC gene expression profiles. **e,** Representative brightfield images of BFP- OSKM- and SRSF1- expressing fibroblasts 1- and 3-days post transgene induction. **f,** Normalized transcriptomic age of SRSF1 induction in M55 across three timepoints for three different induction times. Data (n=2) shows only 2- and 3-day induction maintains long-term transcriptomic reprogramming.

Given the variability in the effects of our perturbations, we used a Uniform Manifold Approximation and Projection (UMAP) to study the differences in the transcriptomes between our 270 cell lines, and we observed distinct clusters corresponding to the three NHDF donors (Fig. 3b). When overlaying the transcriptomic age onto the UMAP plot (Extended Data Fig. 5a), we confirmed that the observed differences were not age-related. Since the WT lines clustered together (Fig. 3b), it is likely that the response of each donor line to our screening workflow introduced transcriptomic differences more pronounced than gene expression changes induced by our perturbations. Nonetheless, the line-specific BFP control confirmed that we could still detect the age-related changes in our edited NHDF lines (Extended Data Table 3). Next, we analyzed the WT lines for age-related changes in the 8 processes of our transcriptomic predictor, and we observed four distinct groups defined by different processes becoming dysfunctional (Extended Data Fig. 5b). These results contextualize the different responses of the same perturbation in multiple cell lines and showcase the variability of human aging, in line with a recent study that described four distinct human aging patterns^45^.

Moreover, our UMAP showed that certain perturbations clustered together (KDM6A, NOTCH1, OSKM, and SRSF1), overcoming donor-specific differences (Fig. 3b). We hypothesized that these genes induced robust transcriptomic reprogramming due to their central role in regulating gene expression as transcription factors or chromatin modifiers, thereby overcoming differences in the starting gene regulatory networks. In agreement with this hypothesis, we observed a negative correlation (Pearson’s r=-0.486; p-value=1.39e-06) between the number of DEGs induced by a perturbation and the mean distance in UMAP space across overexpression lines of the same gene (Fig. 3c). This suggests that the effect of weak perturbations is dependent on the initial cell state (Extended Data Fig. 5c), while perturbations that have a more pronounced effect on gene expression induce a consistent transcriptomic state (Extended Data Fig. 5d-f).

To further study the transcriptomic reprogramming induced by these strong perturbations, we analyzed their effect on cell identity by performing an enrichment analysis for cell-type specific gene sets. All strong perturbations significantly enriched for only fibroblast and mesenchymal stem cell (MSC) identity, except for OSKM, which also induced ESC and iPSC gene expression signatures (Fig. 3d, Extended Data Fig. 5g). Cell morphology confirmed the lack of identity loss in SRSF1-expressing fibroblasts and showed that OSKM overexpression induced a small cell body with multiple projections (Fig. 3e). Lastly, we analyzed the permanence of our transcriptomic reprogramming effect by assaying the transcriptomic age at 0, 6- and 12-days after the end of the transgene induction period. Surprisingly, the data showed that 2- and 3- day inductions maintained only a 10%-15% age reduction over a 12-day period (Fig. 3f), indicating that further dosage and timing optimization could increase the efficiency and durability of age reversal (Extended Data Fig. 5h). As expected, both the OSKM and SRSF1 expressing lines maintained their fibroblast identity after the end of transgene induction (Extended Data Fig 5i).

### Rejuvenating cells using SRSF1 induction

With SRSF1 displaying robust transcriptomic age reversal in all six tested lines (Fig. 3a), and its expression showing a significant decrease during fibroblast aging (Extended Data Fig. 9a), we focused on characterizing its rejuvenation effects in multiple systems with different assays. First, we validated the results from our flow cytometry assays in five new lines, observing a similar >50% decrease in SA-ßGal activity and a ∼40% reduction in proteasome activity (similar to OSKM and opposite to KAT7) (Fig. 4a, Extended Data Fig. 6a, b). Next, we tested the rejuvenation effect of SRSF1 mRNA in more complex fibroblast functions known to decline with age, such as collagen production and response to ROS stress. We observed that a transient 3-day SRSF1 mRNA treatment induced a significant ∼25% increase in collagen production in the five assayed lines and a 30% decrease in the ROS levels upon H_2_O_2_ (Fig. 4b, Extended Data Fig. 6c-f, 7).

**Figure 4.**
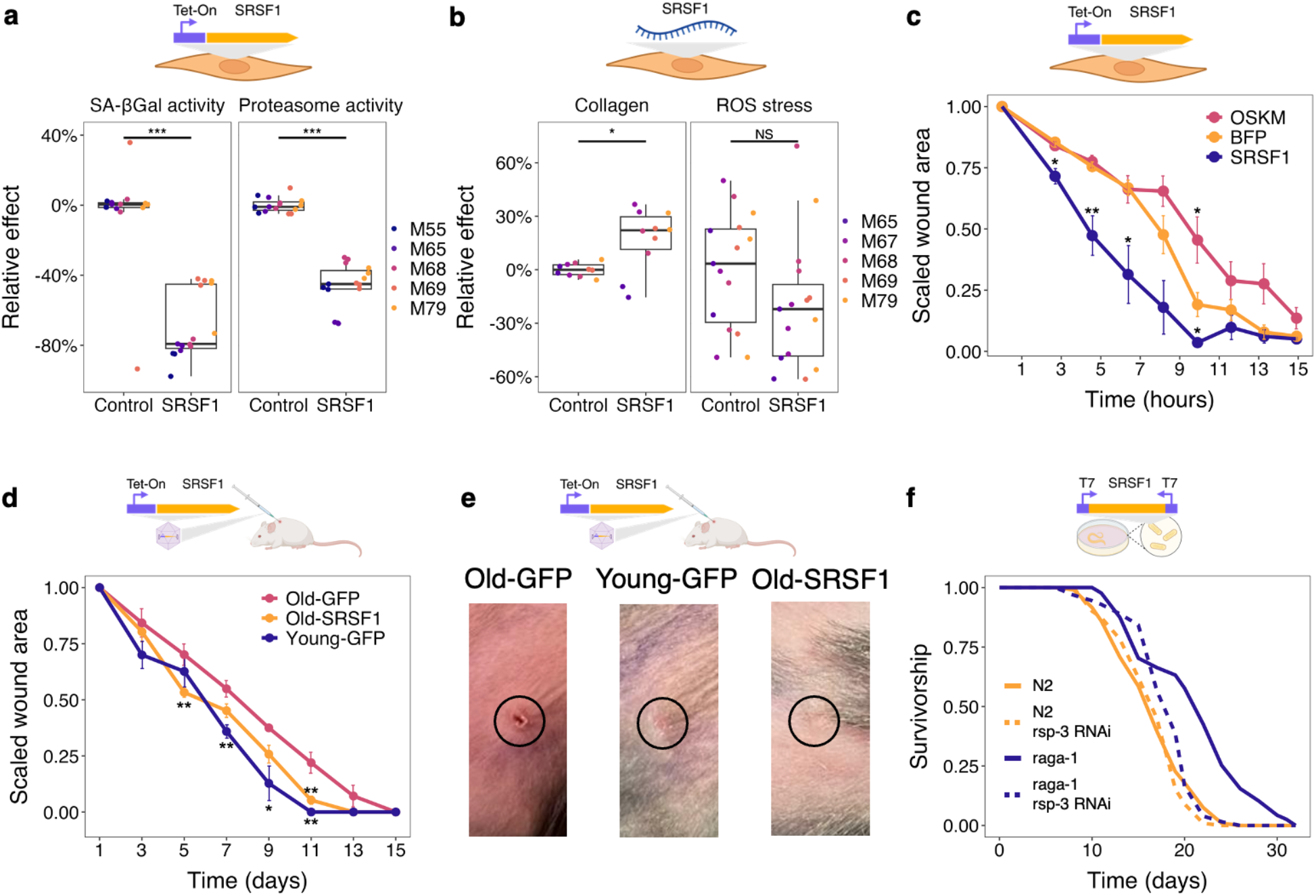
SRSF1 expression induces cellular rejuvenation *in vitro* and *in vivo* a, SA-βGal (**left**) and proteasome (**right**) activity in five differently aged NHDFs (n=3) upon SRSF1 induction shows a significant relative reduction in senescence of ∼80% (p-value = 1.57e-07) and proteasome burden of ∼40% (p-value = 1.53e-10). Data is normalized with respect to the line specific BFP control. **b,** Collagen (**left**) and ROS (**right**) staining in five differently aged NHDFs (n=2) upon SRSF1 mRNA transfection shows a significant relative increase in collagen production of ∼30% (p-value = 0.0133) and a decrease in ROS stress of ∼30% (p-value = 0.127). Data is normalized with respect to the line specific Lipofectamine control. **c,** Scratch assay time course, expressed as wound area scaled to the 0 h time point, for the M79 line after the induction of OKSM, SRSF1, or BFP. Compared to the BFP control, we observed a faster wound area closure in the SRSF1 treated cells and a slower response in those that were induced with OSKM. Data is averaged over replicates (n=3-5), error bars represent standard error of the mean, and significance with respect to the BFP control. **d,** Wound healing time course, expressed as wound area scaled to the day 1 time point, for Young-GFP (n=2), Old-GFP(n=2) and Old-SRSF1(n=4) mice. Compared to the Old-GFP control, we observed a faster wound area closure in the SRSF1 treated mice. **e,** Representative images of skin wounds at day 11 for mice treated with AAV-Tet-GFP or AAV-Tet-SRSF1. **f,** Worm lifespan assay of WT (N2) a long-lived (raga- 1) strains exposed to rsp-3 (SRSF1 homolog) RNAi shows that rsp-3 is required for raga-1 longevity phenotype and has no effect on WT lifespan (Extended Data Table 4). Significance was tested using the two-tailed Student’s t-test and two-way ANOVA.

To further assess the rejuvenation of SRSF1 induction, we tested wound healing efficacy, a key function of dermal fibroblasts, using an *in vitro* scratch assay (Extended Data Fig. 8a). While we did not observe an age-related difference in the migration of WT fibroblasts in our cell culture system (Extended Data Fig. 8b), our data showed that after a 3d induction period, SRSF1- expressing cells reached the midpoint of scratch closure 50% faster than the BFP control, while OSKM induction slightly delayed wound closure (Fig. 4c, Extended Data Fig. 8c), in agreement with previous reports^46^.

Motivated by these results, we tested the effect of SRSF1 gene therapy on mouse wound healing, using intradermal AAV delivery of a Tet-inducible cassette (Extended Data Fig. 9b-d). We observed a large age-related difference in wound healing dynamics, with both the WT and AAV-GFP arms showing a 2-day difference in 50% wound closure and a 4–6-day difference in total healing time between young and old mice (Extended Data Fig. 9e). SRSF1 induction drastically improved the wound healing of old mice, reversing the age-related delay in 50% wound closure and reducing the total healing time to almost that of young mice (Fig. 4d, e). Interestingly, the wound closure phenotype was independent of pre-treatment (induction before wounding), but the healing time reduction required SRSF1 induction throughout the entire process (Extended Data Fig. 9f). Conversely, systemic high-level overexpression of SRSF1 for 21 days did not improve muscle regeneration in old mice (Extended Data Figure 9g), but it also did not induce any observable toxicity or teratomas.

Given that SRSF1 influences translation and has been previously linked to splicing of mTOR targets^47^, we tested whether the *C. elegans* homolog of SRSF1, *rsp-3*, is involved in mTOR longevity. The *raga-1(ok386)* mutant has decreased TORC1 activity and exhibits a 30% increased lifespan compared to WT. Our data showed that while *rsp-3* knockdown has no effect in WT worms, it fully suppressed lifespan extension in the *raga-1(ok386)* mutant (Fig. 4f, Extended Data Fig. 9h, Extended Data Table 4), linking SRSF1 to organismal longevity through the mTOR pathway. Adult *rsp-3* induction experiments could not be conducted due to the transgene activation stalling worm development at the egg stage.

Since SRSF1 is part of the spliceosome, we performed a differential splicing analysis in our fibroblast overexpression lines and observed several alternative transcripts for factors regulating gene expression (Extended Data Table 5), with histone methylation and large ribosomal subunit organization being common across all six lines (Extended Data Fig. 10a). These results agree with the transcriptomic data, which shows a rejuvenation of the histone methylation and translation initiation processes in the SRSF1 overexpression lines (Extended Data Fig. 10b). Moreover, several of the alternatively spliced genes were shown to directly interact with SRSF1^48^, further linking SRSF1 to gene expression changes in histone methylation and translation initiation. Therefore, we propose a mechanism whereby SRSF1 overexpression leads to differential splicing of important regulators of histone methylation and protein translation that reprogram the transcriptome of old cells by inducing youthful gene expression profiles in these processes (Extended Data Fig. 10c).

## Discussion

In this study, we present a method for using the transcriptome in combination with functional assays as *in vitro* aging readouts in large-scale rejuvenation screens. Applying this approach to NHDFs, we discovered nine novel age-modulating genes in human fibroblasts, several of which were also confirmed by our flow-cytometry based assays. The discrepancy in the performance of our hits between the different assays underlines the importance of using multimodal interrogation of the aging phenotype. We believe that such approaches will be instrumental in discovering novel perturbations for reprogramming cell states in the context of aging and other complex phenotypes. Moreover, we showed that both the initial cell state as well as the identity, dosage, and timing of the perturbation influence transcriptomic reprogramming. The data suggested that transcription factors and chromatin modifiers have the most robust effects across different cell states, but can also induce non-specific gene expression changes, necessitating careful dosage optimization.

This observation poses an interesting problem for the development of universal age reversal therapies, which would require strong perturbations without unintended transcriptional shifts that could alter cell identity and function. OSKM induction is a known potent perturbation that has robust rejuvenation effects at the expense of cell identity, thereby being unsafe for therapeutic usage. Here, we discovered SRSF1 expression as a new strong perturbation that can induce robust rejuvenation without de-differentiation or any observable detrimental effects upon systemic overexpression *in vivo*. It is likely that the observed favorable safety profile of SRSF1 is due to the regulation of its own activity through self-splicing^49^. This aligns with our *in vivo* data showing that moderate induction of SRSF1 is beneficial, but high overexpression levels have no observable benefits or toxicity, likely due to the factor’s negative feedback loop. Given that SRSF1 is endogenously expressed in most cells and is transiently activated upon regeneration signals^50^, we believe that the splicing factor can be used as a safe and effective target for fibroblast age reversal. In conclusion, we envision similar transcriptomic reprogramming screens to uncover cell-type specific rejuvenation targets that will serve as the basis for future rejuvenation gene therapies.

## Methods

### RNA-Seq data processing and analysis

Raw RNA-Seq reads were aligned to the GRCh38 human genome using STAR v2.5.2b^51^ and checked for quality control using FastQC v0.11.5^52^. The alignment files were then indexed using SAMtools v1.3.1^53^ and mapped reads were counted using featureCounts from the Subread v2.0.1 package^54^. To reduce the variation between our samples caused by disproportionate sequencing depth, we downsampled the fastq files to 20M reads using Seqtk-1.3^55^. For the overexpression analysis, to address the high number of discarded multi-mapped reads due to the sequence similarity between the endogenous genes and our barcoded genes (Extended Data Table 6), we aligned our reads to a genome index that contains our transgene sequences (including their barcodes) using kallisto v.0.46.2^56^, and we used the tximport^57^ package to generate TPM from estimated counts. We confirmed sample identity using barcode search on the downsampled fastq files with the agrep^58^ tool for approximate string matching with 3 mismatches.

### Training of transcriptomic aging assay

To identify cellular processes predictive of chronological age, we used the Molecular Biology of the Cell (MBotC) Ontology^59^ and empirically selected the third layer to create a set of processes that are potentially informative of aging. In order to quantify the predictability of chronological age from these processes, we built “weak” age predictors for each process as follows: (1) Subset the gene expression matrix to genes in the process under consideration; (2) Performed principal component analysis (PCA) on the subset matrix and replaced it with a matrix containing all principal components. PCA was performed using the implementation in the h2o v3.36.0.2 R package^60^ and imbalanced contribution of individual genes was alleviated by standardizing the data through the ‘transform’ parameter; (3) Trained a generalized linear model (GLM) with 5-fold cross-validation using the ‘h2o.glm’ function in the h2o R package. In particular, the model was based on Gaussian distributions with an identity link function, which standardizes the input before training and implements automatic lambda search with ridge regression; (4) Trained a “strong” age predictor based on the in-bag predictions of all “weak” predictors. The “strong” age predictor is a GLM with 10-fold cross-validation based on Gaussian distributions with identity link function and standardization of the input. However, in contrast to the “weak” models, the strong predictor employed Lasso regression to select only a minimal set of processes that are most informative of chronological age. The combination of processes with non-zero coefficients was considered to be predictive of age. All age predictors were trained on transformed chronological age, using the following piecewise, approximately linear transformation:

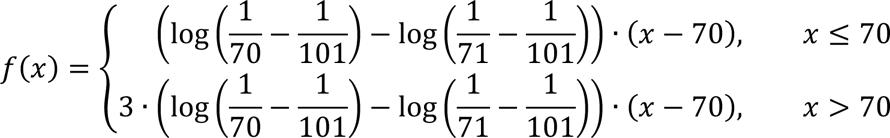

The function is a suitable transformation, since its inverse has the following closed form:

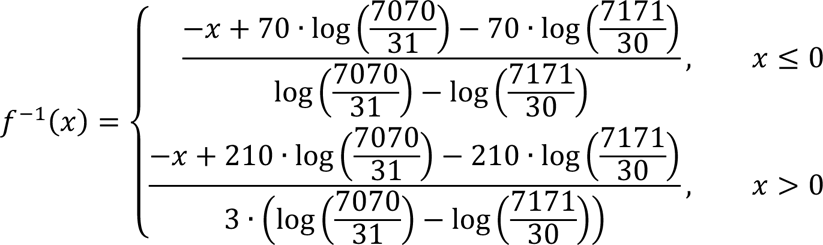

The final transcriptomic aging assay was trained on a set of cellular processes that are predictive of age, using a two-step process. First, for each process, the most predictive set of genes was selected by starting from a model with only an intercept, iteratively adding or removing a gene and scoring each iteration using Akaike’s Information Criterion^61^ by performing linear regression between the current gene set and sample age. In total, 1000 iterations were performed for each process. Second, a GLM, implemented in the h2o v3.36.0.2 R package^60^, was trained on all selected genes of all processes that were selected in the first step. More specifically, the GLM employed Gaussian distributions with an identity link function, lambda search and ridge regression on the standardized gene expression data. Since the number of genes selected in the first step may be larger than the number of training samples, an upper bound on the number of active predictors in the GLM was set to the number of training samples.

### Predicting the Age of Non-training Samples

To remove the variability in raw RNA-Seq data related to sequencing depth, library preparation and other experimental confounding factors, we employed a common pre-processing pipeline across all analyzed datasets. First, training and non-training samples (Extnded Data Table 7) were merged into one matrix and TMM^62^ normalized using the ‘tmm’-function of the NOIseq v2.34.0 R package^63^. Secondly, in case of significant batch effects visible in the first two principal components, batch correction was applied between the training and non-training samples. For the majority of datasets in this work, we employed the naïve Removal of Unwanted Variance (RUV)^64^ algorithm from the RUVnormalize v1.24.0 R package. RUV requires the selection of an appropriate set of control genes that are expected to be uncorrelated to the variable of interest (i.e. age). Thus, we selected appropriate controls as having a low correlation with age in the training data (Pearson’s r < 0.01). Nevertheless, depending on the observed batch effects, other dataset-specific control genes should be included (see individual scripts for the batch correction methods and parameters employed for each dataset). After normalization and batch correction, the transcriptomic predictor was trained on the training data as described in the previous section. The age of non-training samples was then predicted using the ‘h2o.predict’ function in the H2O R package^60^ on the trained model.

### DGEA and cell line comparisons

Differential gene expression analysis was performed using the DESeq2 R package v1.30.1^65^. In particular, DESeq data objects were constructed from raw, untransformed read counts and a design formula reflecting the attempted comparisons. In particular, to detect differentially expressed genes in response to perturbation effects within the screening data, samples of each line were separated and individually processed. Differentially expressed genes were obtained by first running the “DESeq” function with standard parameters followed by the “contrast” function to obtain differentially expressed genes between overexpressed genes and wildtype or blue fluorescent protein (BFP)-transduced controls. For each gene, computed p-values were corrected by applying Bonferroni correction and significance was determined at the 1% level. Cell line identity genes within the large-scale screening assay in three cell lines as well as the validation assay composed of six lines were identified by the following steps. First, all pairwise differentially expressed genes between wild type lines are obtained, as described before. Second, a matrix composed of log2 fold changes across cell lines and all differentially expressed genes was constructed. Next, for each gene in all cell lines, the average log2 fold change between the cell line under consideration and all other lines was computed. Finally, identity genes of a cell line are defined as having the lowest or highest average log2 fold change compared to the other lines. For comparing different cell lines, PCA or UMAP^66^ dimensionality reduction methods were used.

### Cell type signature enrichment analysis

To perform cell type enrichment, we first obtained marker genes for all available cell types from CellMarker database^67^. To increase the accuracy of the obtained markers, we only retained those genes that are either widely used or that were obtained from reviews, and we excluded markers from transformed cells. We then distinguished two cases: (1) the loss of identity compared to a reference set of fibroblast cells and (2) the gain of identity compared to a reference set of induced pluripotent stem cells. For identity loss (1), we sampled a uniform distribution of the training data for our transcriptomic predictor, transformed the gene counts for the selected samples into TPM, and computed the arithmetic mean and standard deviation of each gene. Next, we computed TPM values of the counts for the analyzed sample and subsequently transformed them to z-scores using the previously computed means and standard deviations. Finally, we applied single-sample gene set enrichment analysis implemented in the corto package to obtain significantly lost cell type identities^68^. Since the screening data contains multiple replicates of the same up-regulated genes, we aggregated the obtained p-values using the Sidak method implemented in the aggregation R package^69^. Finally, adjusted p-values were calculated by performing Benjamini-Hochberg correction, with a significant threshold of 0.001. For gained identities (2), we followed the same procedure as described before but employed a reference set of induced pluripotent stem cells for calculating the z-scores of the samples in our genetic screen.

### Computing process activity score

Using the estimated coefficients of our transcriptomic predictor, we quantified the activity of all eight age-associated processes in a sample by computing the scalar product of the model coefficients and the expression values of the corresponding genes. In case of multiple replicates, the activity scores of the same processes in different samples were aggregated into their arithmetic mean. Since RNA-Seq data has been shown to be sensitive to the preprocessing pipeline employed to transform raw read counts, we computed a reference process activity for all samples of our training data. These reference activities define a range of values for each process that correspond to physiological aging and are employed to uniformly scale the activity scores of new samples. Due to the generalized linear model underlying our predictor, lower process activity scores correspond to a younger profile whereas higher values correspond to older transcriptional age.

### Differential Splicing Analysis

Differential splicing analysis was performed using SUPPA2^70^. We first invoked the ‘generateEvents’ subcommand of SUPPA on the GENCODE V29^71^ annotation and parameter setting ‘-e SE SS MX RI FL -f ioe’ to generate all potential splicing events. Next, we measured the proportion spliced-in (PSI) for all potential events by invoking the ‘psiPerEvent’ subcommand on our transcript-aligned, large-scale screening assay (including 6 NHDF lines with SRSF1 over- expression) RNA-Seq data in TPM format. Finally, we invoked the ‘diffSplice’ command with an empirical model (‘-m empirical’) to compare changes between WT and SRSF1 over-expressing lines. Automatic gene correction of p-values is applied by supplying the ‘-gc’ parameter.

### Generating list of candidate age-modulating genes

Candidate target gene list was generated using a network scoring algorithm adapted from the Mogrify method^72^, with modifications for application to complex phenotypes like aging. DGEA was performed on fibroblast RNA-Seq data^2^ using DESeq2 R package v1.30.1^65^ and the associated log-transformed fold change, p-value, and gene expression correlation were combined into a DEG score^31^. To generate a ranked list of factors, we calculated a network score using a weighted sum of DEGscores over a local gene network constructed from STRING^73^.

### Screening library construction

Q5 high-fidelity 2X master mix (NEB M0492) was used to amplify all of the open reading frames (ORFs) from their original vectors (Addgene or ORFeome) in order to add attB sites, the Kozak consensus sequence “GCCACC”, and the WT STOP codon. The amplified fragments were gel purified (QIAGEN 28506) and shuttled into pDONR221 (ThermoFisher 12536017) using the BP Clonase II enzyme mix (ThermoFisher 11789020). The reactions were transformed into 5-alpha competent *E. coli* (NEB C2987H), clones were picked and sequence confirmed using Sanger sequencing. The resulting plasmids were miniprepped (NEB T1010L) and reacted with a barcoded pool of destination vectors (PB-CT3G-ERP2-MG-BC) in a MegaGate reaction^74^. After transforming into 5-alpha cells, clones were picked, sequence confirmed, and barcodes were assigned to specific ORFs (Extended Data Table 6). Final plasmids were miniprepped and used for nucleofection.

### Tissue culture

NHDF lines (Extended Data Table 3) were obtained from the Aging Cell Culture Repository (NIA, Coriell Institute for Medical Studies) and cultured at 37°C, 5% CO_2_, 5% O_2_ in fibroblast media (FM): low glucose DMEM (ThermoFisher 11885-084) supplemented with 15% FBS (GenClone 25-550) and 1% Penicillin-Streptomycin (ThermoFisher 15140122). Before inducing with Doxycycline, cells were switched to media made with Tet System Approved FBS (Takara 631367) to reduce background induction. Media was changed every other day. For generating the overexpression lines, 200,000 cells were nucleofected with 50 fmol transposon and 50 fmol transposase (Super piggyBac Transposase - SystemBio PB210PA-1) or with 300 ng pmaxGFP using the P2 Primary Cell 4D Nucleofector kit (Lonza V4SP-2096) and the Lonza 4D- Nucleofector with the DS-150 program. Cells were recovered at room temperature for 45 min and then plated onto a 24 well plate in 500 μL FM. The following day after nucleofection, dead cells were washed with PBS and media was replenished. 3-4 days post-nucleofection (depending on cell confluency) selection was started using 400 ng/mL Puromycin (ThermoFisher A1113802) in FM (PFM). Cells were selected and expanded at the same time, switching between FM and PFM every 4 days, until the pmaxGFP control reached 0% viability. At the 10 cm dish stage, the cells were frozen in FM with 5% DMSO and stored in liquid nitrogen until thaw. Due to cell line specific sensitivity to the nucleofection and selection process, as well as the increased toxicity of plasmids harboring larger genes, we could not generate overexpression lines for all our 95 genes in all three primary NHDF lines. When assayed, cells were thawed into 2 wells of a 6 well plate in FM and switched to the Tet-free media the next day. 2 days after thaw, cells were harvested, counted, and seeded for RNA-Seq and flow cytometry. For RNA-Seq, each cell line was seeded into 2 wells of a 6 well plate at a density of 60,000 cells/well in FM supplemented with Doxycycline (1 μg/mL for the initial screen and 2 μg/mL for subsequent experiments). For flow cytometry, each line was seeded into 18 wells of a 24 well plate at a density of 10,000 cells/well in FM (the +Dox wells received Doxycycline at 1 μg/mL). 72 h post seeding, the cells were washed with PBS and either stained and harvested for flow cytometry, or lysed using the Monarch DNA/RNA Protection Reagent (NEB T2011) and stored at -80°C.

### RNA Sequencing

RNA was extracted from the cell lysates using the Monarch Total RNA Miniprep kit (NEB T2010) and its quality was spot-checked for random samples using the Bioanalyzer High Sensitivity RNA Kit (Agilent 5067-1513). 100 ng of total RNA was quantified using the Qubit RNA High Sensitivity Assay (ThermoFisher Q32852) and used for library preparation with the NEBNext Ultra II Directional RNA Library Prep Kit for Illumina and the polyA mRNA workflow. The library quality was spot-checked for random samples using the Bioanalyzer High Sensitivity DNA Kit (5067-4626), and then all libraries were quantified using the Qubit dsDNA High Sensitivity Assay (ThermoFisher Q33230) and pooled together. RNA extraction, as well as library preparation, was performed on the same day for all samples that were sequenced together. Sequencing was performed by the Harvard Biopolymers Facility on an Illumina NextSeq or NovaSeq instrument using a 2 × 150 paired-end configuration.

### Flow cytometry

For SA-βGal staining, cells were incubated with 100 nM Bafilomycin A1 (VWR 102513) for 2 h and with 33 nM C_12_FDG (5-Dodecanoylaminofluorescein Di-β-D-Galactopyranoside; ThermoFisher D2893) for 1 h. Mitochondrial membrane potential and proteasome activity were multiplexed by incubating cells with 20 nM TMRM (ThermoFisher M20036), 0.125x proteasome LLVY-R110 substrate, and 0.0625x assay buffer (Millipore Sigma MAK172) for 2 h. Flow cytometry was performed using using either a Cytoflex LX or BD LSRfortessa instrument. Analysis was done using FlowJo (Version 10.8.1).

### IVT and mRNA transfection

The SRSF1 coding sequence was amplified from pAMP531 with the following primers: F 5’- ACTACTTAATACGACTCACTATAGGGATAATGCCACCATGTCGGGAGGTGGTGTG-3’; R 5’-TTTTTTTTTTTTTTTTTTTTTTTTTTTTTTTTTTTTTTTTATGTACGAGAGCGAGATCTG-3’. The PCR product was gel purified using the Monarch DNA Gel Extraction Kit (NEB T2010) and used for IVT reaction using the NEB HiScribe T7 ARCA mRNA Kit (E2065S).

Resulting mRNA was purified using the NEB Monarch RNA cleanup kit (T2040S) and 80 ng of it was transfected into fibroblasts (30,000 cells in 24 well) using 0.75 μL/rxn Lipofectamine MessengerMAX transfection reagent (ThermoFisher LMRNA001).

### Immunofluorescence and imaging

Fibroblasts were fixed using 4% PFA for 15 min at room temeprature (RT) and washed with PBS. Then cells were permeabilized in 0.1% Triton-X in PBS for 15min at RT and washed with PBS. Cells were blocked in blocking buffer (5% FBS, 2% BSA, 0.1% Triton-X-100, 0.05% Tween-20) for 2 h at RT, washed with PBS, and then incubated with Collagen I primary antibody (1:1000 Novusbio NB600-408) in blocking buffer at 4°C overnight. The next day, cells were incubated with secondary antibody cocktail of anti-rabbit Alexa Fluor 488 (1:500, Thermo Fisher Scientific A-11008), 647-conjugated Phalloidin (1:500, Thermo Fisher Scientific A22287), and DAPI (1:50,000, Thermo Fisher Scientific D1306) for 2 h at RT, washed with PBS, covered in ProLong™ Gold Antifade Mountant (Thermo Fisher Scientific P36930) and imaged. For ROS stress quantifications, live cells were first treated with H_2_O_2_ (200 μM, Sigma-Aldrich H1009) for 6 h and washed with PBS. Then, fibroblasts were incubated at 37°C for 45 min with 5 μM CellROX Deep Red oxidative stress reagent (Thermo Fisher Scientific C10422) and 1 μM Calcein AM (Thermo Fisher Scientific, C1430), washed with PBS, and imaged. Z-stack images were captured using a Nikon Ti Spinning disk confocal. Maximum Intensity projections were exported and fluorescent intensity was measured using CellProfiler^75^.

### *In vitro* Scratch Assay

Cells from the M79 line harboring overexpression cassettes for SRSF1, OSKM, or mTagBFP2 were grown in 10 cm dishes and treated with doxycycline (1 μg/mL) for 3 days. A similar protocol was followed for the WT lines, without the addition of doxycycline. Cells were subsequently harvested and seeded into 24 well plates at a density of 120,000 cells per well. The next morning, plates were scratched with a p200 pipette and washed with PBS. Plates were imaged using the Cellcyte live-cell imaging system (Cytena) with a 10x objective at an interval of 1.5 h. Scratch images were stitched, processed, and analyzed as virtual stacks for each time course using Fiji^76,77^. After cropping to the scratch area, the image backgrounds were subtracted using a 10-pixel rolling ball radius and contrast enhanced to 10% saturated pixels, normalized for all images in the stack. The Python package Bowhead v1.1.3^78^ was used to identify the largest contiguous wound area in the images using a threshold of 0.5.

### AAV Production

For *in vivo* wound healing experiments, we chose to use AAV serotype 8 (AAV8) based on previous studies of AAV serotype^79^ infectivity in skin and our own optimization studies, while for the muscle regeneration experiments, we used AAV serotype 9 (AAV9).

HEK293T (ATCC) cells were cultured and maintained in T150 flasks using DMEM high glucose with GlutaMAX (Gibco 10566016), supplemented with 10% FBS (GenClone 25-550) or FetalClone II Serum (Fisher Scientific SH3006603) and 1% Penicillin-Streptomycin (ThermoFisher 15140122). ITR plasmids were transformed into NEB Stable Competent E. coli (NEB C3040H), grown in LB with 100 µg/mL Carbenicillin at 30°C for 18 h, and isolated using the QIAGEN Plasmid Plus Midi Kit (QIAGEN 12943). All other plasmids for making AAV (capsid and helper) were grown at 37°C for 18 h. Plasmids were checked for recombination and mutations using whole-plasmid sequencing. HEK293T cells were transfected in 5-layer flasks at 70-80% confluency. Plasmids were transfected at a ratio of 200:200:100 µg (helper, capsid, ITR plasmid) using PEI. On day 3 following transfection, additional media was added to each flask. On day 6, cells were harvested via dissociation using 5 M NaCl and incubation for 3 h. Dissociated cells were poured into collection bottles and incubated at 4°C overnight to allow sedimentation of cellular debris and nucleic acids. Supernatant was then filtered using 0.22 µM bottle top vacuum filtration systems. PEG8000 was added to the clarified supernatant at a final concentration of 8% (w/v) and placed back at 4°C overnight to allow precipitation. The following day, the mixture was poured into 500 mL conical centrifuge tubes and centrifuged at 3,500g for 15 min. Pellet was then resuspended in 8 mL PBS and 0.8 µL Benzonase endonuclease (Millipore 101695) was added followed by incubation at 37°C for 45 min to remove residual DNA. After incubation, AAV capsids were purified via ultracentrifugation using an iodixanol density gradient and fractions were collected drop-wise in Eppendorf microcentrifuge tubes. Fractions were then assayed via SDS PAGE for purity of AAV capsid proteins. Clean fractions were pooled together and polished using 100kD centrifugal retention filters and cryogenic storage solution made of 5% sorbitol and 0.001% Pluronic F68 in PBS. Purified and polished samples were then quantified using qPCR to obtain a genome copy titer.

### Mice

Young (Stock No: 000664, Sex: Female, Age: 6-8 weeks) and geriatric mice (Stock No: 000664, Sex: Female, Age: 80-90 weeks) were obtained from the Jackson Laboratory (Bar Harbor, ME). Mice were housed in the animal facility of the Harvard Center for Comparative Medicine in Boston, MA according to IACUC guidelines. Except during transgene induction, mice were fed a control diet (Bio-Serv #S4207). For experiments, same-age group mice were housed together and used for control and treatment groups. All animal experiments were approved by IACUC. During surgical procedures, mice were anesthetized with isoflurane. Mouse weights were measured every 2 days until sacrifice to monitor health. No statistical methods were used to predetermine sample size. The mice were randomly assigned to experimental groups, and the investigators were blinded to animal allocation.

### *In Vivo* Wound Healing Experiments

First, each planned wound area was shaved and marked with permanent marker. Next, AAV delivery of our transgene to the mouse skin was achieved through a 100 µL intradermal injection of 1x10^12^ viral genomes per injection for inducible expression (Tet-on) and 1x10^11^ viral genomes per injection for constitutive expression (CAG) into each planned wound area while mice were anesthetized. Marker was reapplied to the planned wound area as needed until wounding. To induce transgene expression in wound healing experiments, mice were fed diets containing doxycycline (200 mg/kg, Bio-Serv #S3888) immediately after injection. Transgene expression was induced for 20 days until wounding. Each mouse received two circular full-thickness excision wounds on the dorsum by a 6-mm biopsy punch. Immediately after mice awoke from anesthesia, they received extended-release buprenorphine (Wildlife Pharmaceuticals) for pain management. Wounds were left uncovered and representative images were taken with a digital camera. Every two days, wound area was measured along the longest and shortest diameter using calipers, which were averaged to obtain the radius and calculate area (A = πr^2^). Wound closure was calculated as the area at each time point divided by the area on day 1. After wounds fully healed, skin was excised for assaying using a 12-mm biopsy punch.

### qRT-PCR

Skin tissue from mice was excised and stored at -80°C. After tissue disruption with a homogenizer (QIAGEN 85600), total RNA was extracted (QIAGEN 74704) and quantified using the Qubit RNA High Sensitivity Assay (ThermoFisher Q32852). Low-quality RNA samples based on low RNA integrity number (RIN) were excluded from downstream analysis. cDNA was synthesized from 100 ng total RNA (NEB #E3010). Real-time polymerase chain reaction (PCR) was performed using Luna® Universal qPCR Master Mix (NEB M3003L) and the Bio-Rad CFX96 Touch Real-Time PCR Detection System (Bio-Rad 1845097). *Gapdh* was used as the endogenous control. Data were analyzed using the comparative CT method. Primers were as follows: *Gapdh* (F 5’-ACTCCACTCACGGCAAATTC-3’; R 5’-TCTCCATGGTGGTGAAGACA-3’) *Srsf1* (F 5’-GCTACGACTACGACGGCTAC-3’; R 5’-AACCACTCTGTTCTCGGACC-3’).

### Muscle regeneration

Cryo-injuries were conducted by exposing the tibialis anterior (TA) muscle and pressing a cold (cooled in dry-ice) flat metal rod to the muscle for 10 seconds. After 7 days, mice were euthanized and regenerating tibialis anterior (TA) muscles were isolated from mice, embedded in O.C.T., and frozen in isopentane cooled with liquid nitrogen for approximately 30 seconds. Frozen TAs were stored at -80°C. Muscle was cryo-sectioned at 10-15 depths covering the middle two-thirds of the muscle (mid-belly) with a 12 μm thickness per section. Frozen tissue sections were then washed with PBS and incubated with anti-Laminin antibody (1:200, Sigma-Aldrich L9393) at 4°C overnight. The next day, sections were washed with PBS, incubated with secondary antibody cocktail of anti-rabbit Alexa Fluor 647 (1:250, Thermo Fisher Scientific A-31573), and DAPI (1:50,000, Thermo Fisher Scientific D1306) for 1 h at RT, washed with PBS, covered in ProLong™ Glass Antifade Mountant (Thermo Fisher Scientific P36980), and imaged on a Zeiss Axio Observer Z1 to evaluate the cross-sectional area (CSA) of regenerating (centrally-nucleated) muscle fibers. CSA of regenerating fibers were analyzed using ImageJ (Fiji) and a custom CellProfiler script. In short, fibers were measured and filtered for those that had a cell nucleus located in either the geometric center or, at a minimum, 1 nucleus length aways from the fiber’s inner edge.

### Worm strains

WBM499 (*raga-1*(*ok386*)II) was made by obtaining VC222 (*raga-1*(*ok386*)II) from the Caenorhabditis Genetic Center, funded by the NIH Office of Research Infrastructure Programs (P40 OD010440), and outcrossing it 6X to in house N2 Bristol wild-type strain.

### Worm lifespans

Lifespan experiments were performed on nematode growth media (NGM) containing 100 μg/ml carbenicillin at 20 °C. Worms were synchronized by timed egg lays using gravid adults. When the progeny reached adulthood (∼72 hours), 120 worms were transferred to fresh plates at 20 worms per plate and this was considered time = 0. Worms were transferred to fresh bacterial lawns every other day until the first deaths (day 10). Survival was scored every 1–2 days, and a worm was deemed dead when unresponsive to 3 taps on the head and tail. Worms were censored due to contamination on the plate, leaving the plate, eggs hatching inside the adult or loss of vulval integrity during reproduction. GraphPad Prism 6 was used to determine median lifespan. Survival curves were compared, and P values calculated using the log-rank (Mantel–Cox) analysis method. Complete lifespan data are available in Extended Data Table 4.

### RNA interference

*rsp-3* RNAi construct came from the Ahringer RNAi library. RNAi experiments were carried out using *E.coli* HT115 bacteria. HT115 bacteria expressing RNAi constructs were grown overnight in LB supplemented with 100 μg/mL carbenicillin and 12.5 μg/mL tetracycline. NGM plus carbenicillin plates were seeded 48 h before use. Respective dsRNA expressing HT115 bacteria were induced by adding 100 μL IPTG (100 mM) one hour before introducing worms to the plate. RNAi was induced from egg hatch. ‘Control’ denotes empty vector HT115 RNAi bacteria.

### Statistical analysis

Statistical analyses were performed with R version 4.0.3, using one- or two-tailed Student’s *t*-tests, ANOVA, or a Z-score. All of the statistical tests performed are indicated in the figure legends. The data are presented as mean or individual points with box plots that show median and quartiles. Error bars represent standard error around the mean.

### Data availability

Accession codes for publicly available data analyzed in this study are provided in Extended Data Table 7. All data generated within this study will be made publicly available before publication.

### Code availability

All original code for computational methods for target identification has been deposited on a Github repository (https://github.com/pranam16/STAMPScreen/), while the source code for transcriptomic assay analyses is available at https://github.com/saschajung/Clock.

## Contributions

A.M.P. and G.M.C. conceived the idea of the study, and A.d.S. conceived the idea of the transcriptomic predictor. A.M.P. wrote the manuscript with input from all co-authors and was involved in all experiments and analyses. S.J. developed the transcriptomic assay with input from A.M.P. and performed computational analyses. H.H.W. contributed to prediction and generation of the candidate list, assisted with experiments and analysis for the RNA-Seq and flow cytometry, performed image data analysis, and assisted with wound healing experiments. F.O. performed the scratch assay, generated AAV, and conducted the wound healing studies. M.S. contributed to the experiments and analysis for the RNA-Seq, flow cytometry, and scratch assay. D.C.B. and Z.K.N performed ROS stress and imaging experiments. M.C.P.M performed worm experiments. N.H. performed muscle regeneration experiments. G.C. and C.V.M assisted with RNA-Seq experiments and image data analysis. G.M.C, A.d.S, W.B.M, and A.J.W supervised the work.

## Acknowledgments

We would like to acknowledge Amanda Graveline, Andyna Vernet, Sarai Bardales, and Melinda Sanchez for assistance with animal work, Isaac Han and Nina Jain for help with AAV production, Yijun Wang for help with library cloning, Nikolaos Dimitrakakis for advice regarding significance testing, and Michael Florea for assistance with muscle injury experiments. We thank Raphaël Ferreira and Sergiy Velychko for comments on the manuscript, as well as Vadim Gladyshev, Ed Boyden, and their lab members for helpful discussions. We thank the BPF Genomics Core Facility at Harvard Medical School for their expertise and instrument availability that supported this work, iHisto for help with tissue sectioning and immunostaining, and New England Biolabs for technical assistance with RNA library preparation. Schematic figures were created with BioRender.com. This work was funded by the Aging and Longevity-Related Research Fund, the Astera Fund at the Silicon Valley Community Foundation, the Wyss Institute Synthetic Biology Platform, and the Spanish Ministry of Science and Innovation (project number: PID2020-118605RB-I00 to S.J.).

---

**Extended Data Figure 1.**
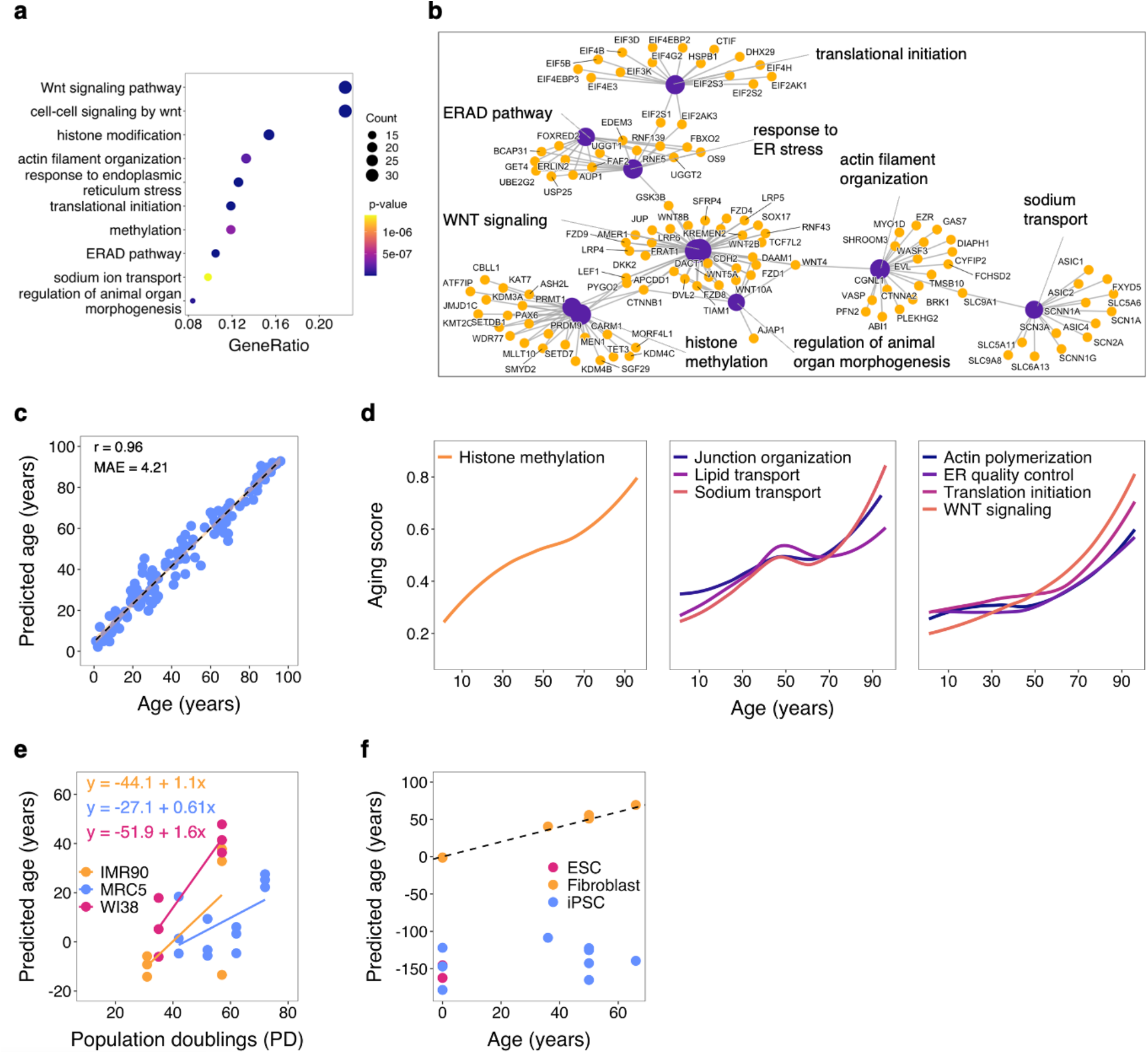
Transcriptomic aging assay correlates with chronological age and passage a, GO term enrichment analysis for selected genes agrees with process annotation and reveals significant enrichment for WNT signaling, histone modification, actin organization, ER stress, translation initiation, sodium transport, and organ morphogenesis. **b,** Network plot of the 146 genes part of the transcriptomic predictor highlights the main processes involved in the aging model, as well as specific genes who have multiple roles in the cell. **c,** Performance of transcriptomic predictor on training data shows high correlation (R^2^=0.96) and low absolute error (MAE=4.2). **d,** Aging score trajectories, calculated as the weighted average of process specific clock genes and normalized across all ages (n=133), for the 8 processes part of the transcriptomic aging assay show 3 distinct clusters: early logarithmic increase (Histone methylation), middle-age inflection (Junction organization, Lipid transport, Sodium transport), and late exponential increase (Translation initiation, Actin polymerization, ER quality control, WNT signaling). **e,** Aging rates of cultured lung fibroblast lines as determined by our transcriptomic predictor. **f,** Predicted ages of NHDFs, age- matched fibroblast derived iPSCs, and embryonic stem cells (ESCs).

**Extended Data Figure 2.**
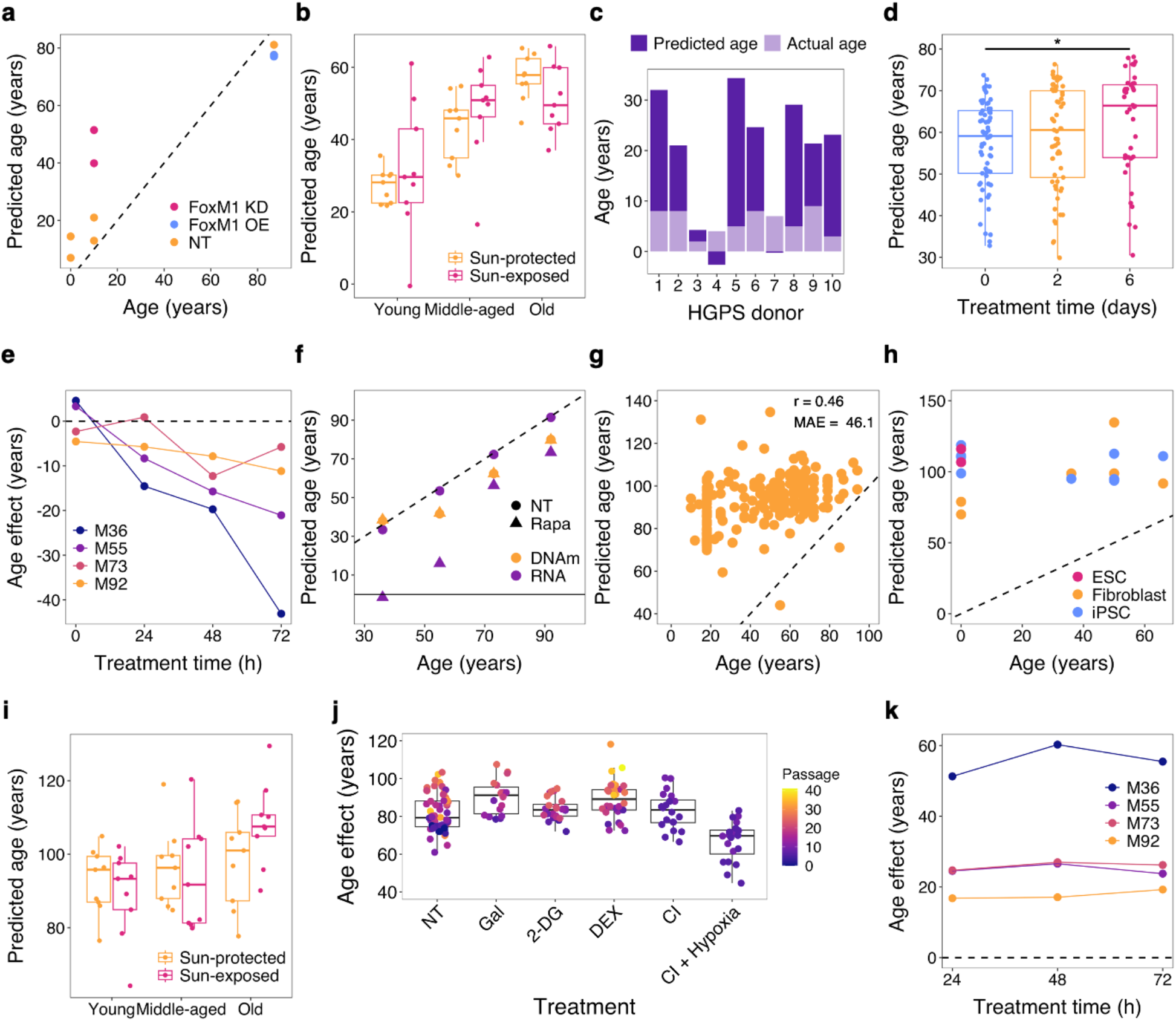
Transcriptomic aging assay responds to anti- and pro-aging treatments a, Predicted ages showing the expected biological effect on NHDFs treated with siRNA for FoxM1 (FoxM1 KD), which was shown to induce senescence, or a constitutively active truncated form of FoxM1 (FoxM1 OE), which was shown to partially rescue aging phenotypes in old cells. **b,** Predicted ages for young, middle-aged, and old NHDFs isolated from a sun protected or sun-exposed area of the body. **c,** Predicted ages of fibroblast samples from Hutchinson-Gilford progeria syndrome (HGPS) donors. **d,** Predicted ages of NHDFs treated with poly(I:C), a known immunostimulant in human cells, for 0, 2 or 6 days. **e,** Age effect (predicted age - age) of Rapamycin (Rapa) (100 nM) as measured by our transcriptomic assay at different treatment timepoints in four different NHDF lines. **f,** Predicted age of Rapa treated samples (100 nM for 72 h) as measured by our transcriptomic assay and the Skin&Blood DNAm clock^5^. **g,** Predictive accuracy of RNAAgeCalc when applied to 12 different published datasets. **h,** Predicted ages of NHDFs, age-matched fibroblast derived iPSCs, and embryonic stem cells (ESCs) using the RNAAgeCalc. **i,** Predicted ages of RNAAgeCalc for young, middle-aged, and old NHDFs isolated from a sun protected or sun-exposed area of the body. **j,** Response of RNAAgeCalc to different *in vitro* stressors. 2-DG = 2-deoxy-D-glucose; DEX = dexamethasone; CI = contact inhibition; hypoxia = 3% O_2_. **k,** Age effect (predicted age - age) of Rapamycin (Rapa) (100 nM) as measured by the RNAAgeCalc at different treatment timepoints in four different NHDF lines.

**Extended Data Figure 3.**
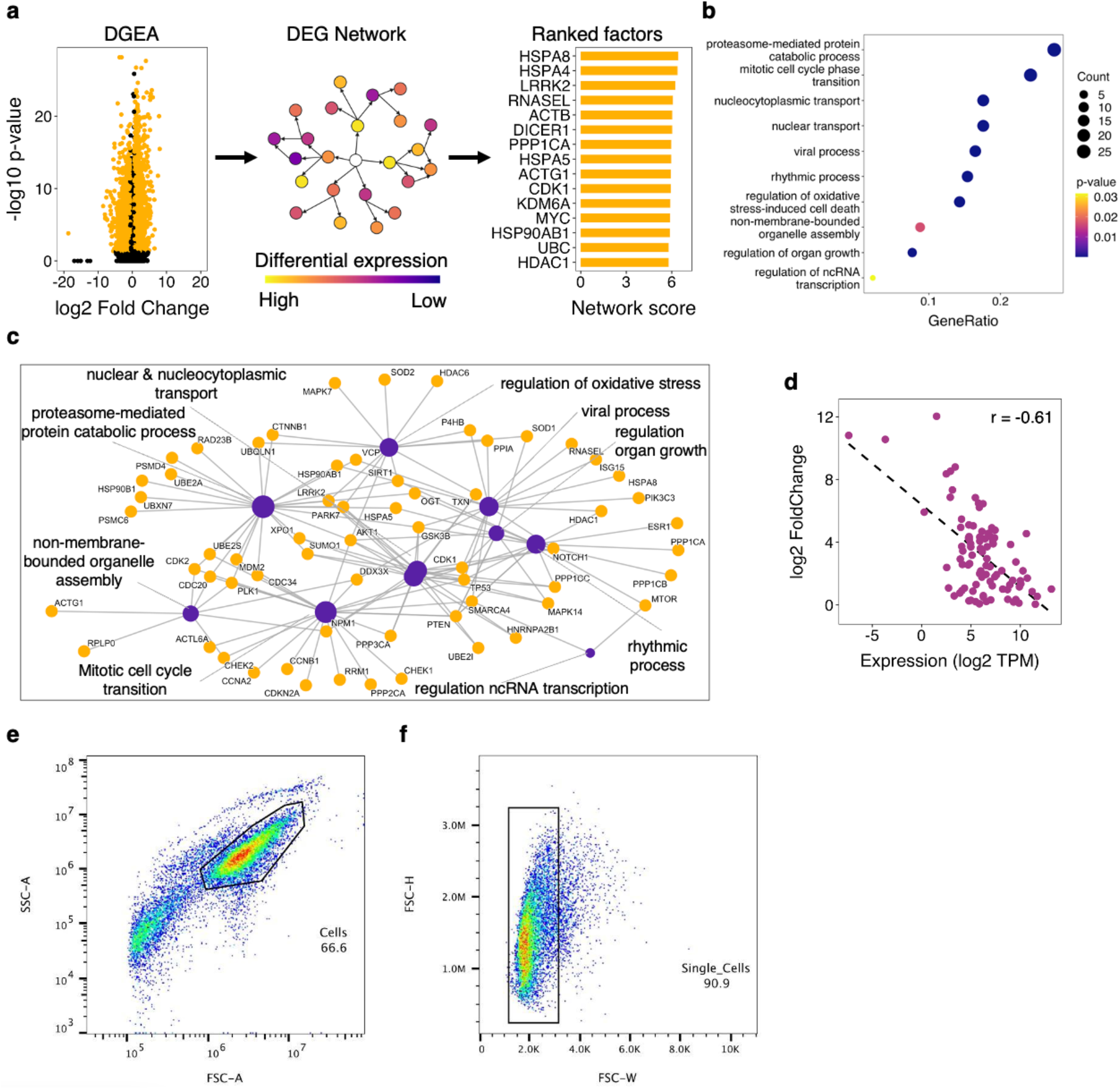
Network scoring algorithm predicts candidate age-modulating genes a, Workflow for predicting candidate age-modulating genes. Differential gene expression analysis (DGEA) was performed on a human dermal fibroblast RNA-Seq dataset generated from 133 individuals with ages 1-94 years old^2^. A gene score (DEGscore) was calculated for each differentially expressed gene (DEG). Then, each candidate gene was assigned a network score based on its local DEG network, derived from the STRING database. Large network scores indicate high connectivity and differential expression levels. **b,** GO term enrichment analysis for candidate aging genes. **c,** Network plot of the 89 candidate genes part of the screen highlights the main processes predicted to influence aging, as well as specific genes who have multiple roles in the cell. **d,** Transgene induction (log2 FoldChange) vs endogenous expression (log2(TPM)) shows a moderate negative correlation of Pearson’s r=-0.61 (p- value = 1.11e-10), with the lowest overexpressed genes having the highest levels of endogenous expression. **e,** Fibroblast cell isolation using forward scatter area and side scatter area, all 18,434 events shown. **f,** Single cell isolation using forward scatter width and height, all 12,277 events shown.

**Extended Data Figure 4.**
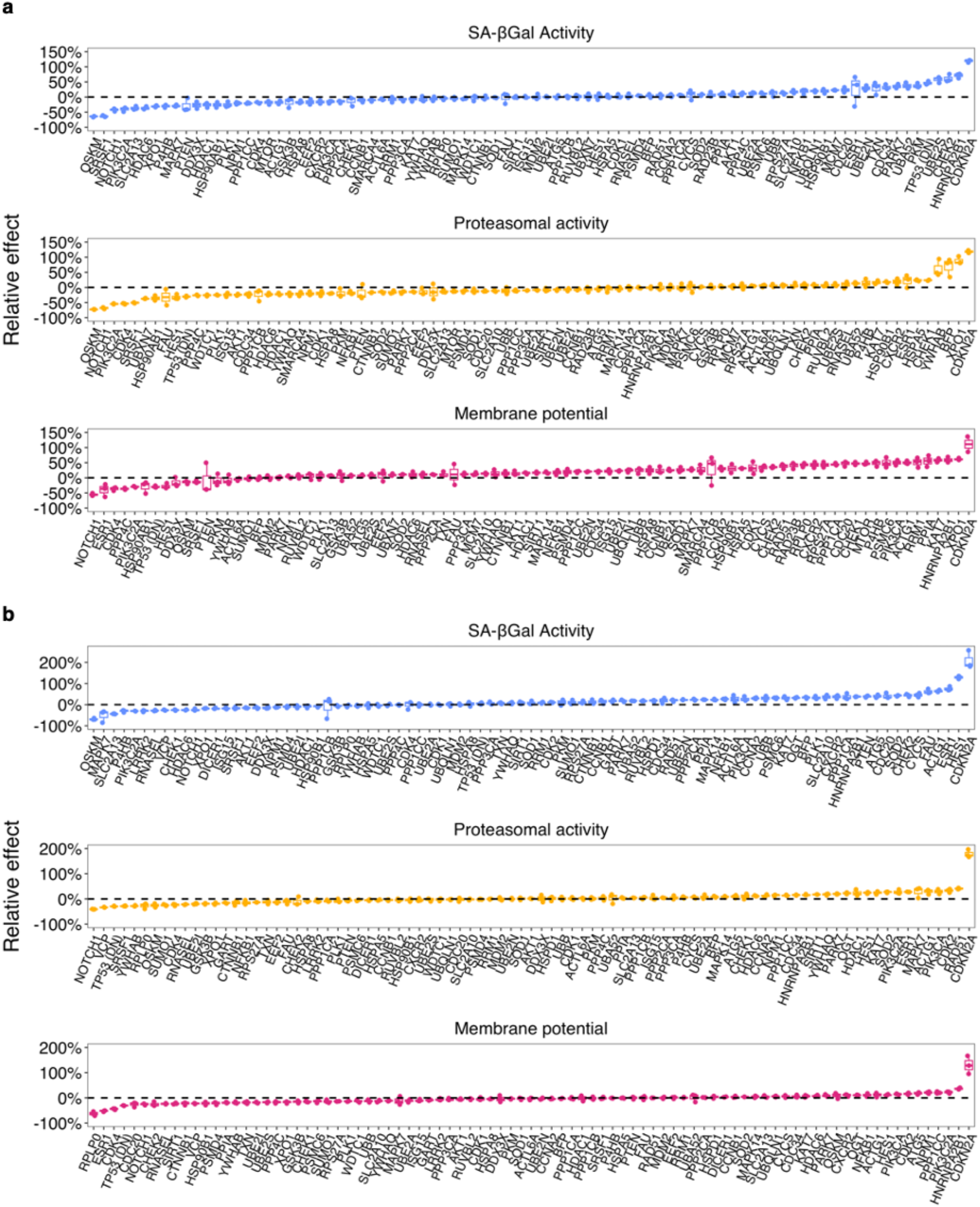
**Staining data for all overexpression lines across the three flow cytometry assays in M55 (a) and M79 (b) fibroblasts Top**. SA-ßGal activity measured for cells expressing gene of interest using C_12_FDG fluorescent substrate. OSKM and SRSF1 reduced cellular senescence, while CDKN2A and CHEK2 increased it. **Middle**. Proteasomal activity measured for cells expressing gene of interest using LLVY fluorescent substrate. NOTCH1 and SRSF1 reduced proteasomal activity while CDKN2A and ESR1 greatly increased it. **Bottom**. Mitochondrial membrane potential measured for cells expressing gene of interest using TMRM dye. NOTCH1 and ESR1 greatly reduced the membrane potential, while CDKN2A and HNRNPA2B1 significantly increased it. Data normalized with respect to the gene specific No Doxycycline control and the line specific BFP control (n=3).

**Extended Data Figure 5.**
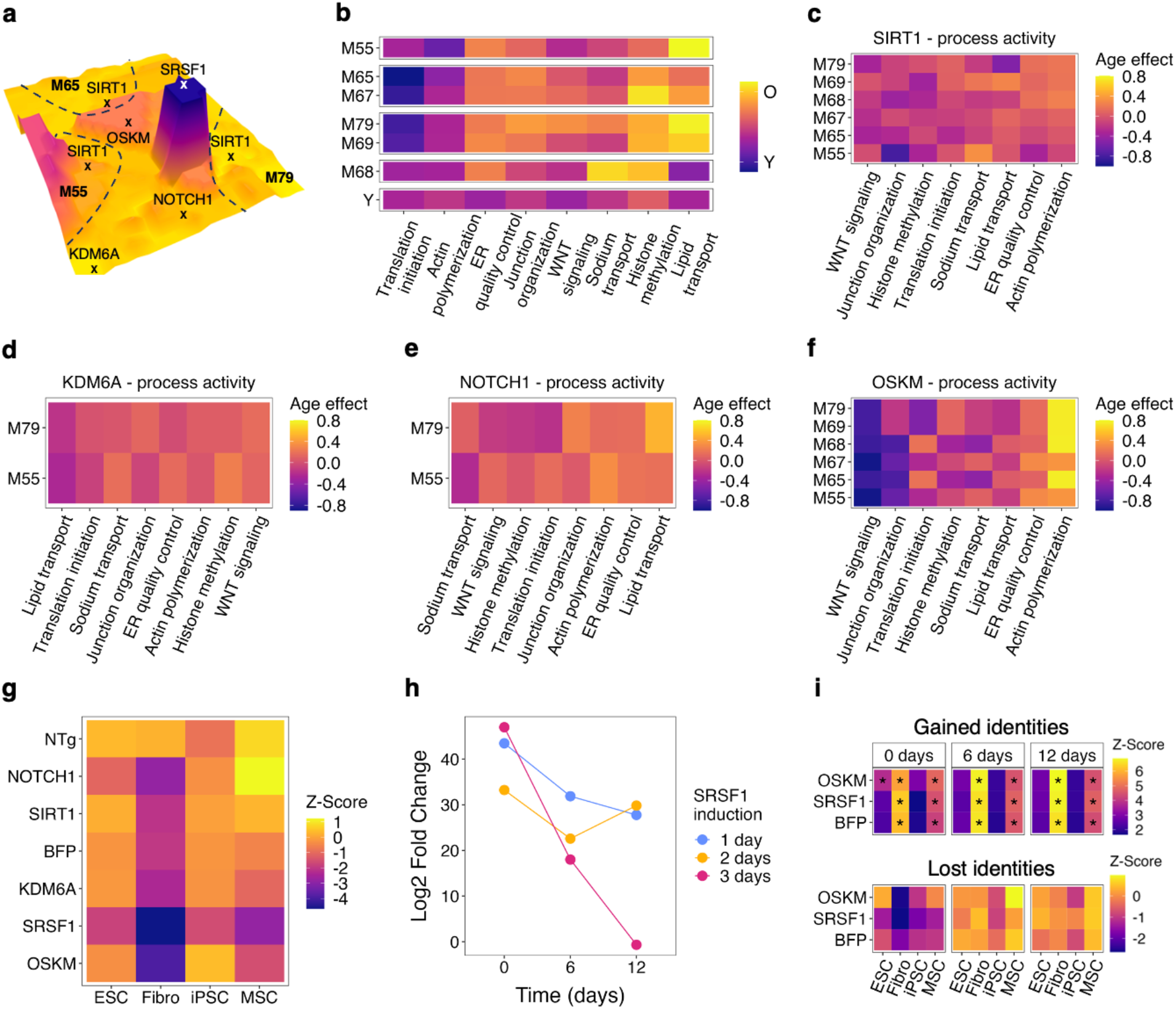
Dynamics of transcriptomic reprogramming across different perturbations a, 3D landscape of all 480 NHDF transcriptomes from our large-scale screen, with the x-y plane representing the UMAP coordinates from Fig. 3b and the z-axis representing transcriptomic age. **b,** Heatmap of differential (older age - younger age) process activity in the 6 NHDFs and an averaged young control. We observed four distinct clusters based on the nature of the process dysfunction: M55 = lipid transport; M65&M67 = histone methylation, M69&M79= lipid transport and histone methylation, M68 = sodium transport and histone methylation. **c,** Scaled age effect of SIRT1 overexpression across six lines (n=2). We observed no consistent effects, but we noticed that the responder lines (M65, M67, M68, M79) all had moderate rejuvenation in WNT signaling. **d,** Scaled age effect of KDM6A overexpression across M55 and M79 (n=2). We observed a weak and consistent pro-aging effect in WNT signaling, histone methylation, and actin polymerization, with a rejuvenation effect in lipid transport. **e,** Scaled age effect of NOTCH1 overexpression across M55 and M79 (n=2). We observed a consistent pro-aging effect in lipid transport, ER quality control, actin polymerization. **f,** Scaled age effect of OSKM overexpression across all 6 lines (n=2). We observed a consistent rejuvenating effect in WNT signaling, junction organization, and translation initiation (except for M65 and M68) and pronounced pro-aging effect in actin polymerization. **g,** Negative enrichment analysis for cell identities with respect to a fibroblast transcriptome. Data is averaged across the 3 NHDF lines (M55, M65, M79) and shows a reduction of fibroblast-specific gene expression in the SRSF1 and OSKM lines, which does not reach statistical significance. **h,** Log2 Fold Change for SRSF1 expression in the M55 transgenic lines across 3 timepoints for 3 different induction times relative to the BFP control. Data (n=2/timepoint) shows SRSF1 expression decreasing after Dox withdrawal, with different dynamics for the three induction protocols. **i,** Cell type signature enrichment analysis for gained (top) and lost (bottom) identities for the 1-day induction time course in M55. Data (n=2) show a normalization of fibroblast gene expression program after the end of transgene induction.

**Extended Data Figure 6.**
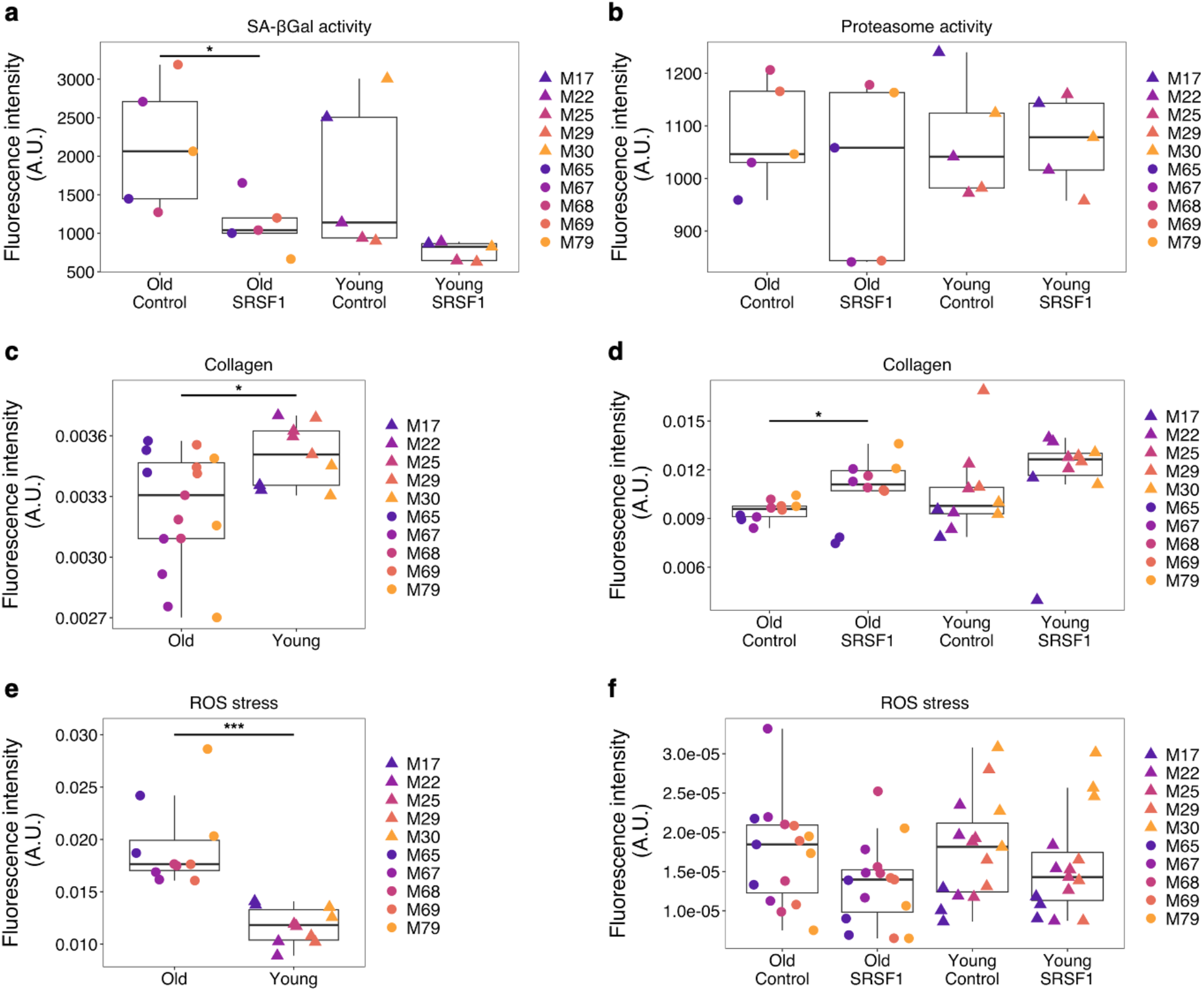
SRSF1 mRNA transfection rejuvenates old fibroblasts a, SA-βGal activity in 5 young and 5 old NHDFs (n=3) transfected with SRSF1 mRNA or Lipofectamine control shows a significant 50% reduction in senescence in the old lines. **b,** Proteasome activity in 5 young and 5 old NHDFs (n=3) transfected with SRSF1 mRNA or Lipofectamine control. **c,** Collagen staining in 5 young and 5 old NHDFs (n=3) shows an 8% decrease in mean collagen production in the old fibroblasts. **d,** Collagen staining in 5 young and 5 old NHDFs (n=3) transfected with SRSF1 mRNA or Lipofectamine control shows a significant 14% increase in mean collagen production in the old treated fibroblasts. **e,** ROS staining in 5 young and 5 old NHDFs (n=2) shows a 39% increase in mean ROS production in the old fibroblasts. **f,** ROS staining in 5 young and 5 old NHDFs (n=3) transfected with SRSF1 mRNA or Lipofectamine control shows a mean relative decrease in ROS stress of ∼22% (fails to reach significance). Significance is tested using the two-tailed paired (for SRSF1 vs control) or unpaired (for young vs old) Student’s t-test.

**Extended Data Figure 7.**
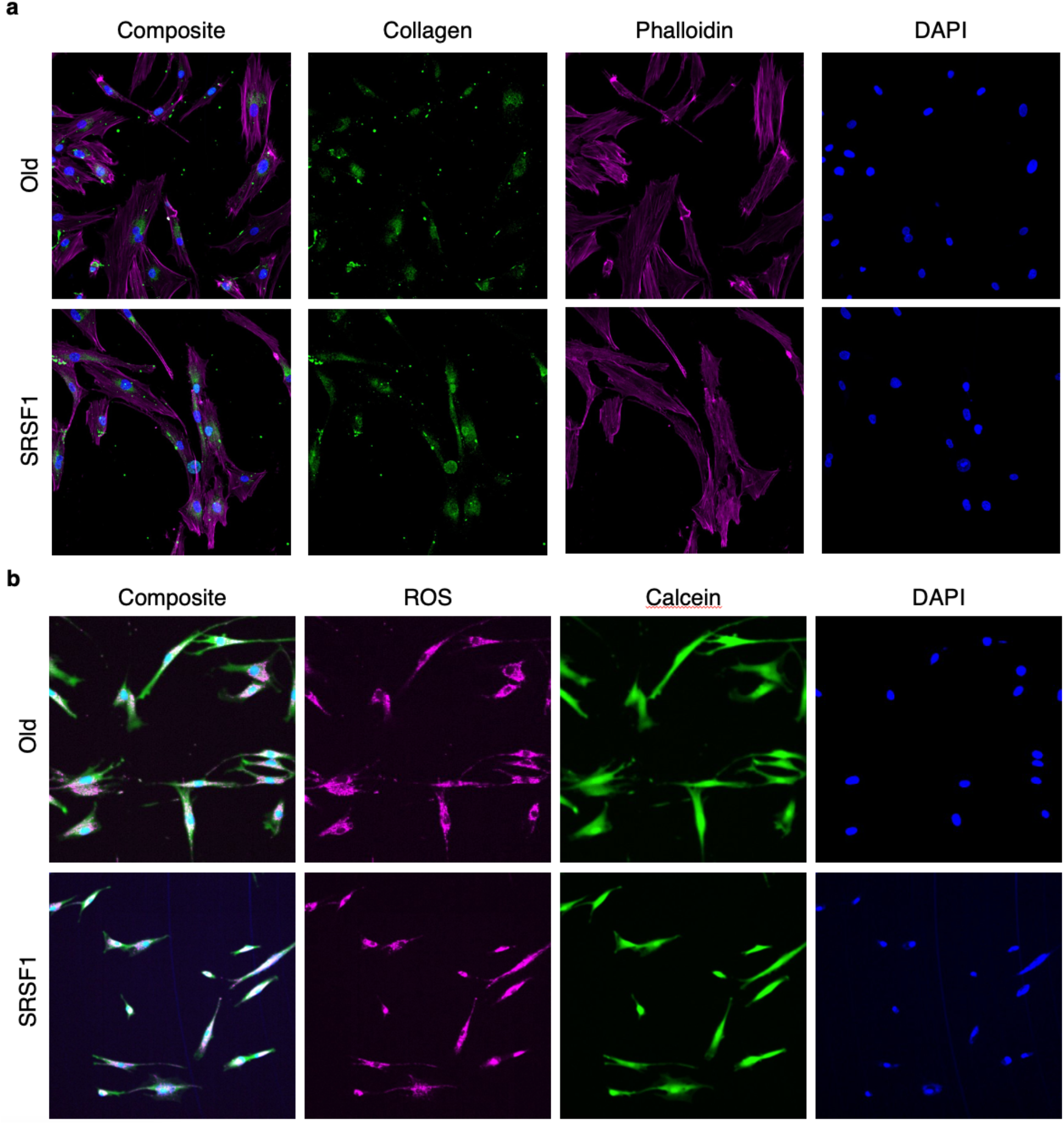
Representative images of oxidative stress and collagen staining in transfected NHDFs a, Confocal images of collagen in lipofectamine control (top) and SRSF1-mRNA transfected fibroblasts (bottom). Collagen (green) was measured within the cell’s actin cytoskeleton (Phalloidin; Deep red). DAPI (blue) was used to localize cells. SRSF1-mRNA transfected cells showed increased Collagen throughout the cell. **b,** Lipofectamine control (top) and SRSF1-mRNA transfected fibroblasts (bottom) were treated with H_2_O_2_ for 6 h before assessing reactive oxygen species (ROS) using CellROX (Deep Red). ROS analysis was quantified in viable cells, stained with Calcein-AM (green) and DAPI (Blue). Both conditions showed accumulation of ROS, however SRSF1- mRNA transfected cells showed decreased ROS intensity diffused throughout the cell.

**Extended Data Figure 8.**
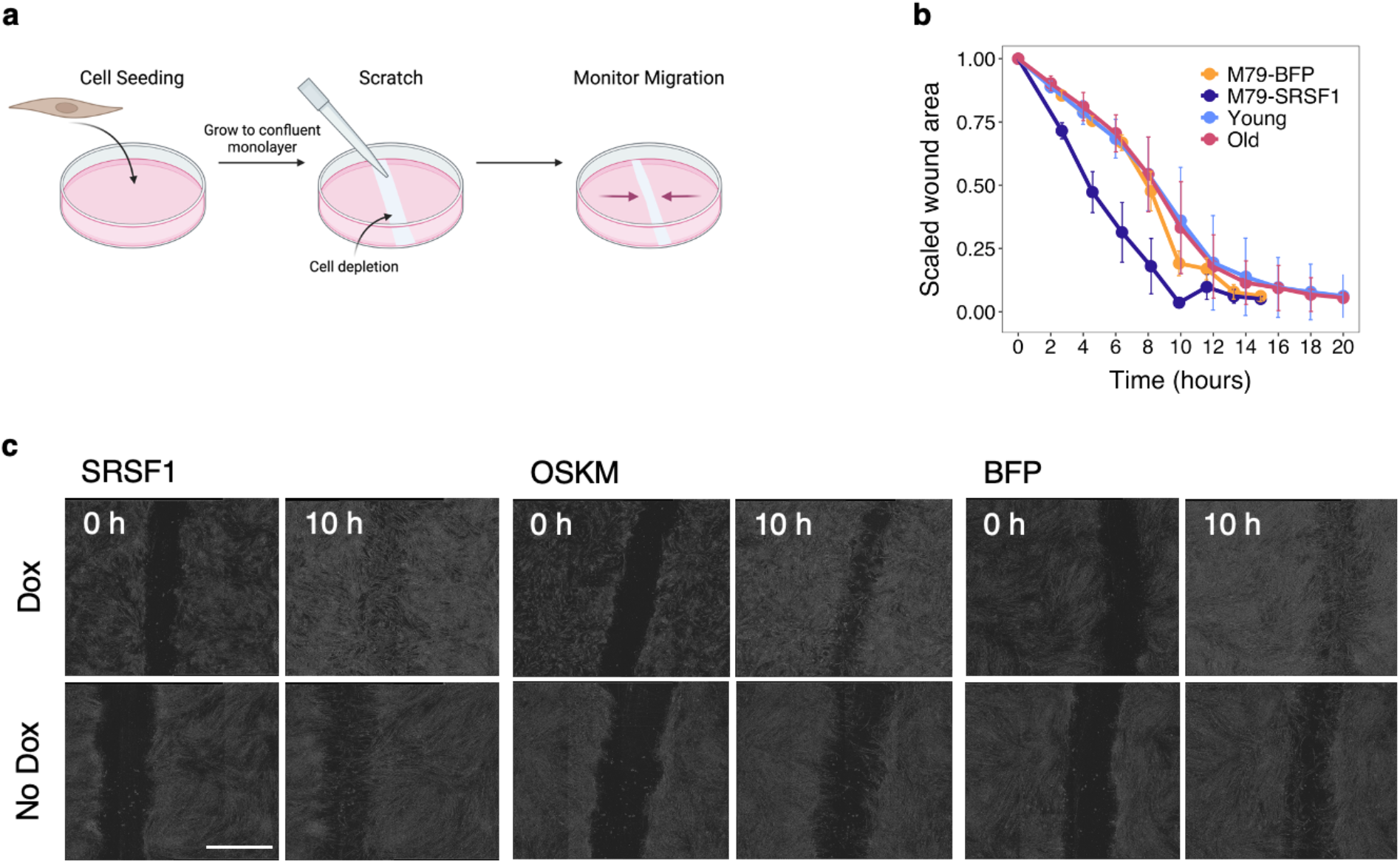
Scratch assay in NHDFs a, Experimental diagram for *in vitro* scratch assay. **b,** Scratch assay time course, expressed as wound area scaled to the 0 h time point, for the M79 line expressing BFP or SRSF1 (n=3-5) and for young and old WT lines (n=3-5 averaged over 5 different NHDF lines per group). **c,** Representative bright-field images of SRSF1-, OSKM-, or BFP-induced cells at 0 h and 10 h after scratch. Scale bar, 1 mm. Experiments were performed independently at least three times with similar results.

**Extended Data Figure 9.**
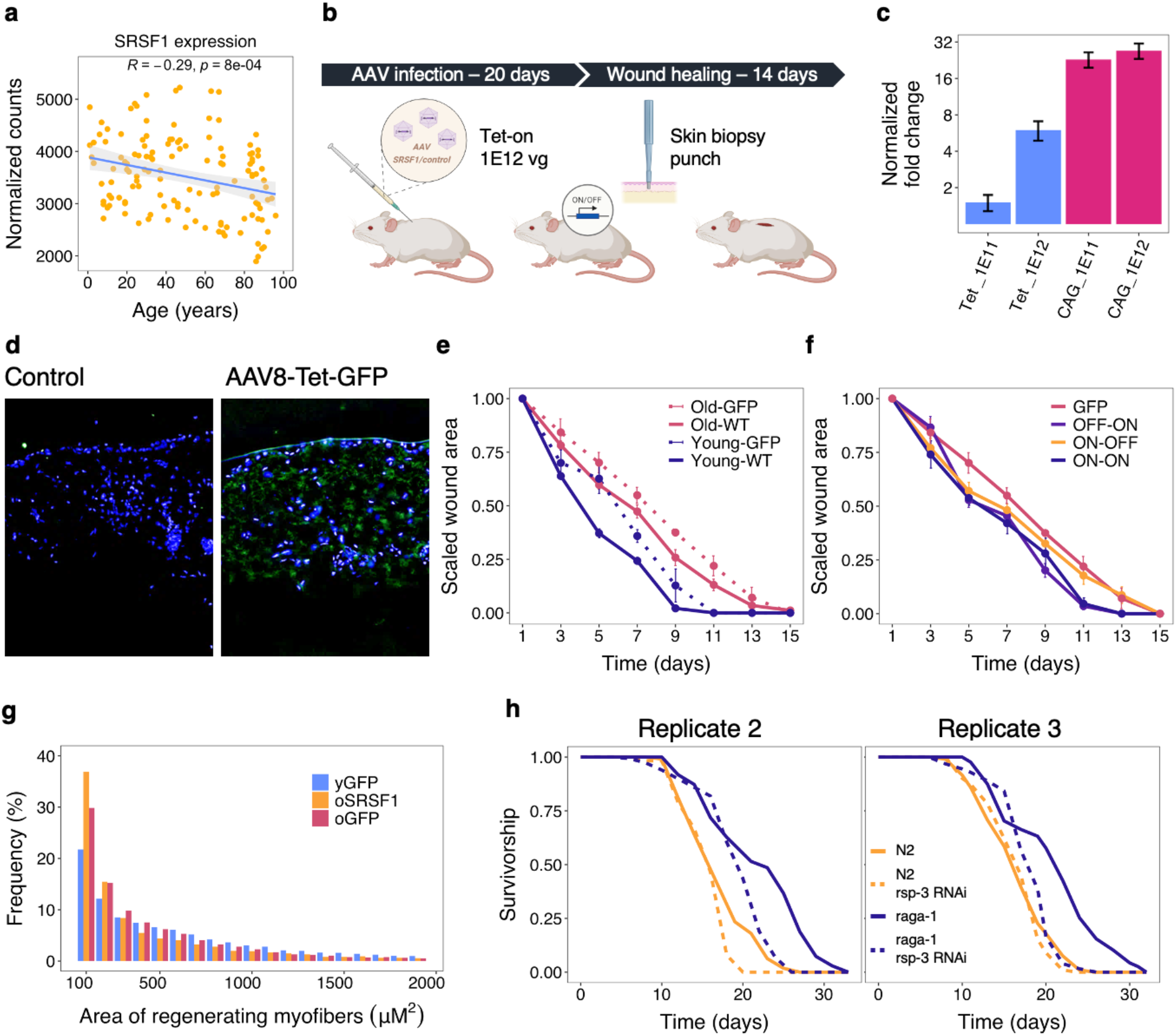
SRSF1 modulation *in vivo* a, Normalized SRSF1 expression in NHDFs of different ages shows a significant decline with age of ∼50% between young and old fibroblasts. **b,** Experimental diagram for *in vivo* wound healing assay. Mice were injected with 1E12 viral genomes of AAV8 carrying either a Dox-inducible SRSF1 or GFP cassette. 20d after injection, mice were wounded in the injection area using a punch biopsy, and the wound healing was monitored for 14 days after. **c,** Normalized fold change of skin SRSF1 expression 21 days after injection with Dox-inducible or constitutive AAV vector at 2 different doses (n=1/group). **d,** Skin immunohistochemistry for GFP in mock-injected and AAV8-Tet-GFP injected mice 21d post-injection. **e,** Scaled wound area of WT (n=8 mice/group) and AAV-GFP (n=2 mice/group) injected mice shows large age-related differences in wound healing dynamics, with the gene therapy slowing down the process equally in both young and old mice. Data is averaged over replicates (n=2 wounds/mouse), and error bars represent standard error of the mean. **f,** Scaled wound area for old mice injected with AAV-Tet-GFP or AAV-Tet- SRSF1 with different induction times (n=2 mice/group). ON-OFF = Dox was withdrawn at the time of wounding. OFF-ON = Dox was added at the time of wounding. ON-ON = Dox was added at the time of injection and kept throughout the wound healing process. Data is averaged over replicates (n=2 wounds/mouse), and error bars represent standard error of the mean. **g,** Regenerating myofibers distribution in mice treated with AAV9-CAG-GFP or AAV9- CAG-SRSF1systemic gene therapy 7 days after cryo-injury. Data shows old mice have a higher proportion of small fibers compared to the young ones, indicating the delayed healing response. SRSF1 did not significantly affect the healing process, likely due to the high overexpression achieved by the CAG promoter. **h,** Two additional replicates for the worm lifespans show complete cancelation of *raga-1* longevity phenotype in *rsp-3* knockdown worms.

**Extended Data Figure 10.**
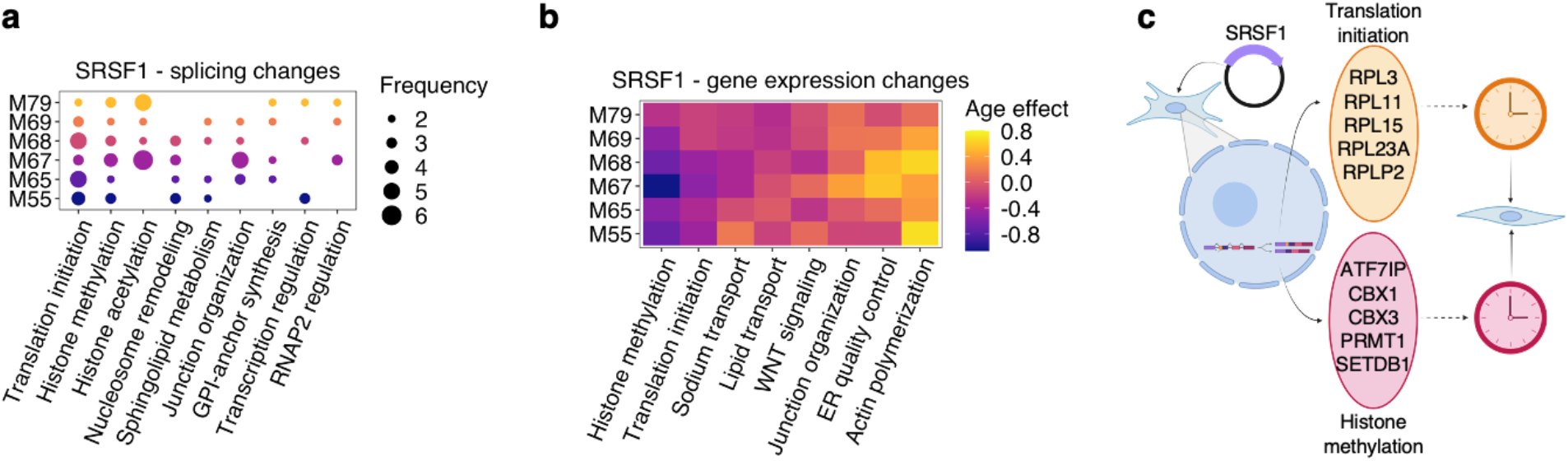
Proposed mechanism for SRSF1 rejuvenation a, Enrichment analysis for differentially spliced genes across six SRSF1-expressing lines (n=2) shows translation and histone methylation as the main affected processes. **b,** Scaled age effect of SRSF1 overexpression on the clock processes across six lines (n=2). Negative age effects represent younger gene expression profiles. We observed high correlation between the process effects across all 6 lines, with a significant improvement in histone methylation and translation initiation. **c**, Proposed mechanism for SRSF1 rejuvenation. The alternative splicing of several genes part of the translation initiation and the histone methylation pathways leads to a beneficial effect on the expression of clock genes part of the same processes, which ultimately induces a younger transcriptome in the treated cells.

